# The Kinship Matrix: Inferring the Kinship Structure of a Population from its Demography

**DOI:** 10.1101/2021.04.12.439517

**Authors:** Christophe F. D. Coste, François Bienvenu, Victor Ronget, Juan-Pablo Ramirez-Loza, Sarah Cubaynes, Samuel Pavard

## Abstract

The familial structure of a population and the relatedness of its individuals are determined by its demography. There is, however, no general method to infer kinship directly from the life-cycle of a structured population. Yet this question is central to fields such as ecology, evolution and conservation, especially in contexts where there is a strong interdependence between familial structure and population dynamics. Here, we give a general formula to compute, from any matrix population model, the expected number of arbitrary kin (sisters, nieces, cousins, etc) of a focal individual *ego*, structured by the class of *ego* and of its kin. Central to our approach are classic but little-used tools known as genealogical matrices. Our method can be used to obtain both individual-based and population-wide metrics of kinship, as we illustrate. It also makes it possible to analyze the sensitivity of the kinship structure to the traits implemented in the model.

## 1 Introduction

The demography of a population determines its level of inbreeding and its kinship structure, i.e. the network of kin relationships between the individuals of the population. Population geneticists have long used non-overlapping generation models to study the effects of demography on kinship structure, for instance to estimate the frequency of consanguineous marriage (Barrai et al., 1962; Hajnal, 1963). Models with overlapping generations have also been considered – from the seminal work of Goodman et al. (1974) to the recent advances by Caswell (2019, 2020) – but they focus on specific kin relationships. Thus, a general method has yet to be developed to infer kinship structure from population dynamics in structured populations, and little is known about the influence of specific life history traits (or vital rates in general) on inbreeding levels, inclusive fitness and kinship/familial structures. Yet those questions are central to fields such as ecology, demography, evolution, genetics and conservation.

In this article, we pave the way towards such a general method by showing how classic tools from the study of matrix population models can be combined in a new way to infer kinship structure from steady-state population dynamics. More specifically, we explain how to compute the expected number of kin of a focal individual *ego*, as a function of the classes of *ego* and of its kin, directly from the projection matrix of a population. Examples of applications include studying how inbreeding depression leads to genetic Allee effects in populations of small effective sizes or modeling the eco-evolutionary demography of social species (where the vital rates of individuals may strongly depend on cooperative and competitive interactions with kin). Readily usable implementations of our methods are available at 10.5281/zenodo.4680715, see SM.1.

### 1.1 A brief history of kinship inference in structured populations

The first demographers to work on the estimation of kin distribution from the demographic rates of a structured population based their computations on age-structured, one-sex populations at the demographic steady state, and focused on a few key kin relationships (Le Bras, 1973; Goodman et al., 1974, discussed in Pavard and Coste, 2020). Their method, still widely used today, consists in computing the distribution of maternal age-at-childbirth from the Euler–Lotka equation (Lotka, 1939), and then use it to derive, for each age-class, the probability that the mother/grandmother of a focal individual is alive, as well as the expected number of daughters, sisters, aunts and nieces of that individual.

Shortly after, mathematicians attacked the problem from the field of branching processes. First, for unstructured populations without overlapping generations, using Bienaymé–Galton–Watson processes (Waugh, 1981; Pullum, 1982); then, by adding age as a category and considering multitype Crump-Mode-Jagers processes (e.g., Nerman and Jagers, 1984). However, the technical nature of these papers and their focus on sometimes abstract results has made them go largely unnoticed by demographers. For instance, despite being directly relevant to the matter, the work by Joffe and Waugh (1985) is only cited nine times in the mathematical literature – and not once in a biology or demography journal.

As of today, the inferred kin frequencies provided by Goodman et al. (1974), which have been in constant use since their publication, are still the state of the art (Pavard and Coste, 2020). These take the form of a limited number of *ad hoc* formulas, each specific to a kin relationship, that only apply to age-structured populations where at most one offspring can be produced at a time. Recently, Caswell (2019, 2020) has put this framework in matrix form and extended it to age×stage models, by giving a system of recursive equations expressing the number of kin at time *t* + 1 as a function of the number of kin a time *t*. However, before the present work computing the expected number of any kin in any life cycle remained an open problem and a necessary step in the development of a general theory of the interplay between kinship and demographic processes (Pavard and Coste, 2020).

### 1.2 Specific challenges to overcome

Inferring the complete kinship structure of an arbitrary structured population poses several challenges. First, in general class-structured models the distribution of the class at birth is not as easily inferred as in age-structured populations, where it is obtained from the Euler–Lotka equation. We solve this problem by using the genealogical Markov chains first introduced by Demetrius to define population entropy (Demetrius, 1974, 1975). These somewhat little-known tools have recently proved useful to tackle various questions (Bienvenu and Legendre, 2015; Bienvenu et al., 2017).

The fact that several offspring can be produced at the same time constitutes another complication, because this prevents segregating genealogies as done in Goodman et al. (1974) and Caswell (2019, 2020). Accounting for the possibility of same-litter sisters requires special care and fine-grained information on reproduction.

Finally, the main difficulty is arguably the generalization of the calculations to arbitrary kin relationships. This requires combining two different timescales: the demographic timescale (where time is expressed in fixed projection intervals, such as one year) and the genealogical one (where time is expressed in number of generations). We do so by working out all the genealogies and associated trajectories of individuals in the life cycle that correspond to a given kin relationship.

### 1.3 The kinship matrices

In order to describe the kinship structure of a population, we need a generic notation for arbitrary kin relationships. In one-sex populations, we can use the following: we say that two individuals (*i*_1_*, i*_2_) are (*g, q*)*-kin* when their most recent common ancestor is separated from *i*_1_ by *g* generations and from *i*_2_ by *q* generations. For instance, the (1, 0)-kin of the focal individual *ego* is its mother, and its (0, 1)-kin are its daughters; similarly, its (1, 1)-kin are its sisters; its (1, 2)-kin its nieces; its (2, 1)-kin its aunts; etc (see Figure 1A). The same convention was used by Atkins (1974) and Pullum (1982), but with the order of *g* and *q* reversed.

**Figure 1:**
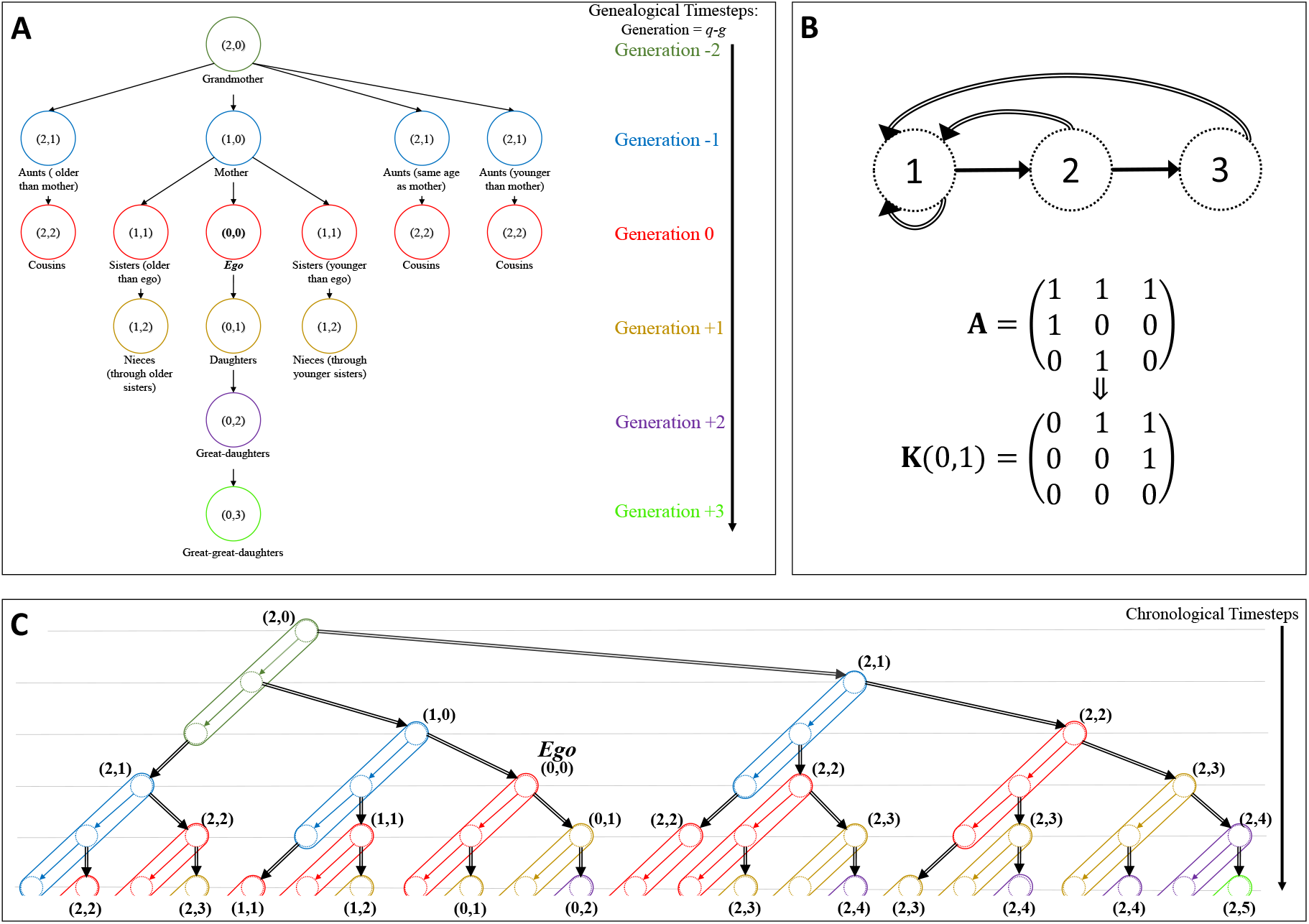
**(A)** Kin relationships, as described in terms of (*g, q*)-kin of the focal individual *ego*. Each line (and colour) corresponds to a generation. To get to the (*g, q*)-kin of *ego*, one has to go “up” *g* generations in order to reach the most recent common ancestor of *ego* and its (*g, q*)-kin; and then “down” *q* generations to reach the (*g, q*)-kin themselves. **(B)** Simple example of an age-structured life-cycle, with the corresponding projection matrix and kinship matrix **K**(0, 1) giving the expected number of daughters of *ego*. In this toy model, all individuals survive to age 3 and then die, and they produce exactly one offspring each year. Thus, individuals aged 2 have exactly one daughter (aged 1) and individuals aged 3 have exactly two daughters (one aged 1 and one aged 2). **(C)** The genealogy of *ego* for the population described in (B), with the corresponding kin structure. Each dotted circle corresponds to an individual at a given age, and each solid oval to an individual’s life trajectory. The colors corresponds to the generation with respect to *ego*, as in (A). Here we have assumed that *ego* was born when its mother was aged 1, and that the mother of *ego* was born when its grandmother was aged 2.

We describe the kinship structure of a class-structured population through the *kinship matrices* **K**(*g, q*), where the (*i, j*)-th entry of **K**(*g, q*) is the expected number of (*g, q*)-kin alive in class *i* of *ego* in class *j*. From the point of view of *ego* – which is what matters for most applications – these matrices completely characterize the (expected) kinship structure of the population. An example of the genealogy of a focal individual, together with the corresponding population model and kinship matrix for daughters, is given in Figures 1B and 1C.

The rest of this article is organized as follows: in Section 2, we explain how to compute the kinship matrices directly from the projection matrix of the population. We then illustrate the method in Box 2 using the life-cycle of the ground squirrel *Spermophilus dauricus* from Luo and Fox (1990); other detailed examples of applications can be found in SM.2. Finally, we conclude by discussing the limitations, implications and possible extensions of our method in Section 3.

## 2 Method

Our method is applicable to populations at the demographic steady state whose dynamics are governed by a matrix population model with a primitive (that is, irreducible and aperiodic) projection matrix **A** = (*a*_*ij*_). It can also be used with some reducible matrices (see SM.3), but here we assume primitivity, for simplicity. Recall that *a*_*ij*_ is the per-capita contribution of an individual in class *j* at time *t* to the abundance of class *i* at time *t*+1. The asymptotic growth rate *λ* is given by the dominant eigenvalue of **A** and the stable class distribution by the corresponding right eigenvector **w** = (*w*_*i*_) such that Σ_*i*_*w*_*i*_ = 1.

To compute the kinship matrix **K**(*g, q*), we decompose the projection matrix **A** into its survival and reproductive components – that is, into **A** = **S** + **F**, where the *survival matrix* **S** = (*s*_*ij*_) is such that *s*_*ij*_ is the probability that an individual of class *j* survives into class *i*, and the *fertility matrix* **F** = (*f*_*ij*_) is such that *f*_*ij*_ is the expected number of offspring of class *i* produced by an individual of class *j*. Mathematically, we need reproduction and survival to be independent (it is otherwise not possible to retrace the life trajectory of an ancestor and then use this trajectory to compute its number of offspring). This assumption is never applicable in post-breeding models, where individuals have to survive a full projection interval before they can reproduce. We therefore recommend using pre-breeding models whenever possible and avoiding post-breeding models with our method. Birth-flow models can also be used as long as the covariance between survival and reproduction is kept in mind as a potentially significant source of discrepancy. Finally, note that one way to deal with covarying reproduction and survival (e.g, due to trade-offs) is to add (survival, fertility) classes to the model.

Finally, to take into account models with several offspring per time-step, we need the *same-litter newborn sisters matrix* **Z** = (*z*_*ij*_), where *z*_*ij*_ is the expected number of same-litter sisters of *ego* that have just been born in class *i*, knowing that *ego* is in class *j*. Unlike **S** and **F**, the matrix **Z** is not always directly available, and to compute it one will typically need to make additional assumptions on the offspring distribution. We give expressions of **Z** under three of the most common scenarios: when at most one offspring is produced by projection interval; when there is only one class of offspring; and when the total number of offspring follows a Poisson distribution and each offspring is independently allocated to a newborn class.

The three matrices **S**, **F** and **Z** are all we need to compute any kinship matrix **K**(*g, q*), as summarized in Box 1. Python and MATLAB implementations of the method are provided (see SM.1). In the rest of this section, we detail the reasoning behind the expression of **K**(*g, q*) and give examples of aggregated metrics of the kinship or relatedness structure of a population that can be derived from it. Concrete examples of application are given in Box 2 and SM.2.

#### Box 1: Expression of the kinship matrix K(*g, q*)

- Input: let **A** = **S** + **F** be the projection matrix of the population, where **S** is the survival matrix and **F** is the fertility matrix. Let **Z** be the same-litter newborn sisters matrix (see Section 2.4).
- Compute the genealogical matrices **P**_**S**_ = (*p*_**S**_(*i, j*)) and **P**_**F**_ = (*p*_**F**_(*i, j*)) defined by

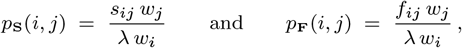

where *λ* is the asymptotic growth rate (dominant eigenvalue of **A**) and **w** the stable class distribution vector (dominant right eigenvector of **A**).
- For all integers *n* ≥ 0 and *ℓ* ≥ 1, let 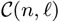 denote the set of vectors **k** = (*k*_1_, …, *k*_ℓ_) of non-negative integers such that *k*_1_ + · · · + *k*_ℓ_ = *n* (when *ℓ* = 0 or *n <* 0, let 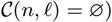.
- For any **k** = (*k*_1_, …, *k*_ℓ_), let 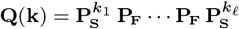 and

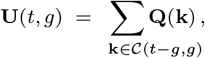

with the convention that **U**(0, 0) = **I**, the identity matrix, and that **U**(*t, g*) = **0** when the sum is empty.
- For any **k** = (*k*_1_, …, *k*_ℓ_), let 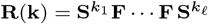 and

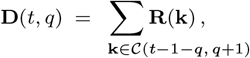

with the convention that **D**(*t, q*) = **0** when the sum is empty.
- The kinship matrix **K**(*g, q*), whose (*i, j*)-th entry is the expected number of (*g, q*)-kin of class *i* of a focal individual of class *j*, is **K**(0, 0) = **I** and, for (*g, q*) ≠ (0, 0),

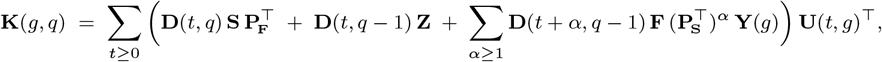

where **Y**(*g*) = **I** if *g* = 0 and 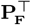 otherwise.

### 2.1 The genealogical Markov chain

The central tool of our approach is the genealogical Markov chain introduced by Demetrius (1974, 1975) to define population entropy. Despite its early introduction in the field of matrix population models, this Markov chain has remained mostly associated with population entropy (see, e.g., Tuljapurkar, 1982, 1993) and has so far failed to become part of the standard toolbox of population ecologists and demographers. However, it has recently proved useful to study questions such as the generation time (Bienvenu and Legendre, 2015) and the optimal aggregation of classes (Bienvenu et al., 2017). Our work provides yet another example of the usefulness of this tool.

The transition matrix of the genealogical Markov chain associated with **A** is the matrix **P** = (*p*_*ij*_), where

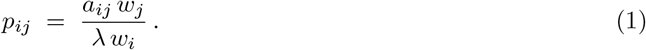

This Markov chain describes the sequence of classes that we encounter as we go “up” the genealogy of the population by following the lineage of an individual, backwards in time. That is, *p*_*ij*_ is the probability that an individual in class *i* at time *t* was either alive and in class *j* at time *t* − 1; or else had its mother in class *j* at that time. Thus, the probability distribution of the class of the ancestor (in the sense of a younger self or a genealogical ancestor), *t* time steps ago, of an individual currently in class *i* is given by **e**_*i*_**P**^*t*^, where *e*_*i*_(*j*) = 1 if *j* = *i* and 0 otherwise (note that the formalism of Markov chain uses right-multiplication to project probability distributions, as in **x**(*t* + 1) = **x**(*t*) **P**, whereas population projection matrices use left-multiplication). See Bienvenu et al. (2017) for more on genealogical Markov chains.

As we did with **A**, we can split the transitions of the genealogical matrix into a survival and a reproductive component. We then have **P** = **P**_**S**_ + **P**_**F**_, where **P**_**S**_ = (*p*_**S**_(*i, j*)) and **P**_**F**_ = (*p*_**F**_(*i, j*)) are given by

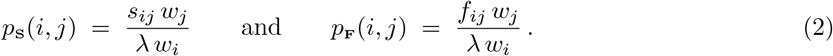

Accordingly, *p*_**S**_(*i, j*) is the probability that an individual in class *i* at time *t* was alive and in class *j* at time *t* − 1, while *p*_**F**_(*i, j*) is the probability that it was born between times *t* − 1 and *t* to a mother in class *j*. Thus, for instance, the (*i, j*)-th entry of **P**_**S**_**P**_**F**_ is the probability that the mother of an individual from class *i* was in class *j* two time-steps ago *and* that the first transition when going up the genealogy was a survival transition, while the second one was a reproductive transition.

We are now able to: (1) project the class of the ancestor of *ego* backwards in time (within generations using **P**_**S**_ and between generations using **P**_**F**_); and (2) project the descent of an ancestor forwards in time (within generations using **S** and between generations using **F**). With this, our strategy to compute the number of (*g, q*)-kin of *ego* is to go “up” the genealogy to locate its *g*-th ancestor; and then “down” to compute the *q*-th descendants of that ancestor. However, to account for same-litter sisters, we also need to look at the (*g* − 1)-th ancestor of *ego*.

### 2.2 Distribution of the (*g* − 1)-th ancestor of *ego*

Let us start by assuming that *g* ≥ 1 (the case *g* = 0 will be treated separately later). For *t* ≥ 1, let **U**(*t, g*) = be the matrix whose (*i, j*)-th entry is the probability that *t* − 1 time steps ago the (*g* − 1)-th ancestor of *ego* was in class *j*, given that *ego* is currently in class *i*. To compute **U**(*t, g*), we need to consider all the ways that *g* − 1 generations can pass in the span of *t* − 1 time steps. For any integers *n* ≥ 0 and *ℓ* ≥ 1, let

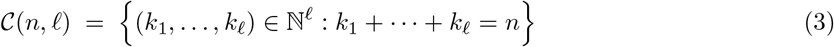

denote the set of vectors of length *ℓ* whose components are non-negative integers that sum to *n* (with 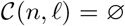 when *ℓ* = 0 or *n <* 0). Such vectors are known as *weak compositions of the integer n into ℓ parts*, and there are efficient algorithms to list them (see SM.4). The weak compositions 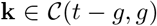 exactly encode the sequences of *t* − 1 “survival” or “reproduction” events that contain *g* − 1 “reproduction”: indeed, **k** = (*k*_1_, …, *k*_*g*_) corresponds to the sequence

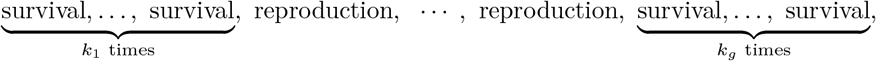

Thus, letting 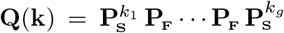 – whose (*i, j*)-th entry is the probability that, starting from class *i*, after *k*_1_ + … + *k*_*g*_ + *g* − 1 steps one ends in class *j* after having encountered *k*_1_ survivals, then one reproduction, then *k*_2_ survivals, etc. – we have

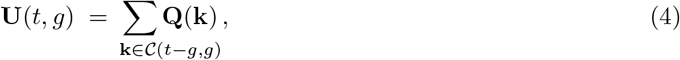

with the convention that **U**(*t, g*) = **0** when the sum is empty (e.g., if *t < g*).

As examples of the formula, note that for *g* = 1 and *t* ≥ 1, 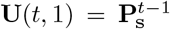, and for *g* = 2 and *t* = 2, 3, 4, …

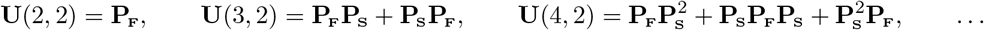

Finally, letting **U**(*t, g*) = (*u*_*ij*_(*t, g*)), the probability, given that *ego* is currently in class *i*, that its *g*-th ancestor gave birth to its (*g* − 1)-th ancestor in class *ℓ*, from class *k* and between times *t* and *t* − 1 before the present is *u*_*iℓ*_(*t, g*) *p*_**F**_(*ℓ, k*). The reason why we need to look at both the (*g* − 1)-th and the *g*-th ancestor of *ego*, rather than at its *g*-th ancestor only, is that we will need this information to account for the (*g, q*)-kin *ego* that are descendants of same-litter sisters of its (*g* − 1)-th ancestor. Note that the probability, given that *ego* is currently in class *i*, that its *g*-th ancestor gave birth to its (*g* − 1)-th ancestor from class *k* and between times *t* and *t* − 1 before the present is the (*i, k*)-th entry of **U**(*t, g*)**P**_**F**_.

### 2.3 Expected number of q-th generation descendants

Let **D**(*t, q*) be the matrix whose (*i, j*)-th entry is the expected number of *q*-th generation descendants in class *i*, after *t* − 1 time steps, of an individual in class *j* (not counting offspring produced before the first time-step). As before, to compute **D**(*t, q*) we need to consider all the ways that *q* generations can pass in the span of *t* − 1 time steps. For **k** = (*k*_1_, …, *k*_ℓ_), let 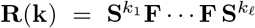. Recall that the expected number of descendants in class *i* of an individual in class *j*, after *t* − 1 time steps (and counting the initial individual) is the (*i, j*)-th entry of the matrix

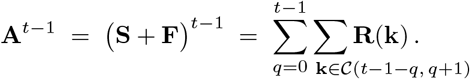

For instance, for *t* − 1 = 3,

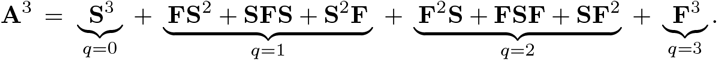

In these expressions, the terms corresponding to a fixed value of *q* correspond to the *q*-th generation descendants of the initial individual. In other words,

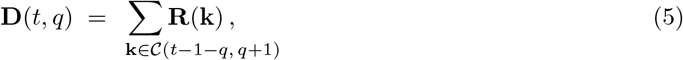

again with the convention that **D**(*t, q*) = **0** when the sum is empty (e.g, when *q <* 0 or *t* − 1 *< q*).

### 2.4 Expected number of (g, q)-kin

Let *ego* be in class *j*, and recall that we assume *g* ≥ 1. For *t* ranging over the positive integers and *k, ℓ* over the classes of the model, the events “the *g*-th ancestor of *ego* gave birth to its (*g* − 1)-th ancestor in class *ℓ*, from class *k*, and *t* − 1 time-steps before the present” form a complete system of events. Each of these events has probability *u*_*jℓ*_(*t, g*) *p*_**F**_(*ℓ, k*). Thus, all we need in order to compute the expected number of (*g, q*)-kin of *ego* is the expected number of *q*-th generation descendants of an individual that was in class *k* at time *t* before the present and had an offspring born in class *ℓ* between times *t* and *t* − 1. For this, we need to distinguish between descendants of younger, older and same-litter sisters of the (*g* − 1)-th ancestor of *ego* (see Figure 2).

**Figure 2:**
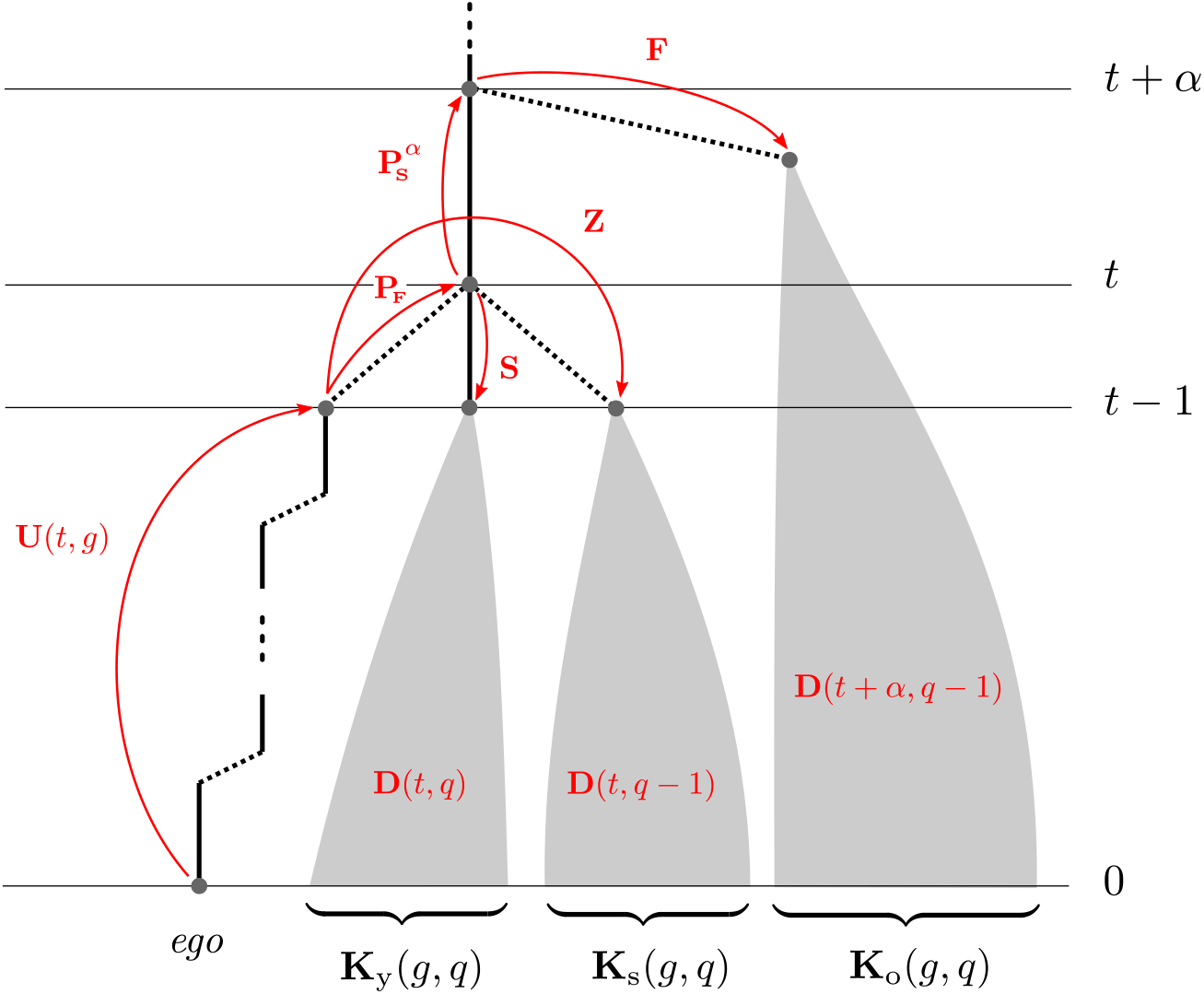
Genealogy of the population, showing the various contributions to the kinship matrix **K**(*g, q*). On the left is the lineage of *ego*, up to its *g*-th ancestor. The solid black lines correspond to survival and the dotted ones to reproduction. On the right are the *q*-th descendants of the *g*-th ancestor of *ego*, grouped according to whether they are descendants of younger, same-litter or older sisters of the (*g* − 1) *th* ancestor of *ego*. The weights of the red arrows have to be multiplied along a path to get the corresponding contribution to the expected number of (*g, q*)-kin (matrix transposes have been omitted for simplicity).

#### Descendants of younger sisters of the (*g* − 1)-th ancestor

Between times *t* and *t* − 1 before the present, the *g*-th ancestor of *ego* survives from class *k* to class *m* with probability *s*_*mk*_. From there, the expected number of *q*-th generation descendants that it leaves in class *i* after the remaining *t* − 1 time steps is *d*_*im*_(*t, g*), the (*i, m*)-th entry of the matrix **D**(*t, g*). The expected number of class-*i*(*g, q*)-kin of *ego* through younger sisters of its (*g* − 1)-th ancestor is therefore Σ_*t*≥0_ Σ_*k,ℓ,m*_ *d*_*im*_(*t, q*) *s*_*mk*_ *u*_*jℓ*_(*t, g*) *p*_**F**_(*ℓ, k*) – which, in matrix notation, is the (*i, j*)-th entry of

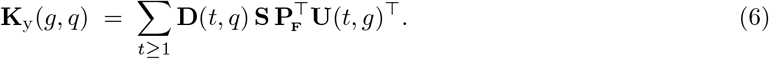

#### Descendants of older sisters of the (*g* − 1)-th ancestor

To compute the expected number of descendants of older sisters of the (*g* − 1)-th ancestor of *ego*, we have to look at the *g*-th ancestor at times *t* + *α* before the present, for each *α* ≥ 1, to see how many older sisters were born then and how many (*q* − 1)-th generation descendants each of these left. Using the same reasoning as before, we get:

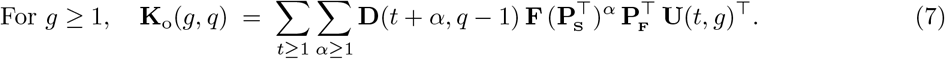

#### Descendants of same-litter sisters of the (*g* − 1)-th ancestor

Unlike for descendants of older and younger sisters, for which knowing about the *g*-th ancestor of *ego* was sufficient, here we also need to know about its (*g* − 1)-th ancestor. This is because knowing that the *g*-th ancestor gave birth to the (*g* − 1)-th one *t* time-steps before the present biases its number of offspring for this reproductive event. To understand this bias, we have to know the class in which the (*g* − 1)-th ancestor was born. Indeed, if we sample an individual uniformly at random in class *i* and then look at its mother, then the mother is sampled proportionally to its number of offspring in class *i*. Let *F*_*ij*_ have the distribution of the number of offspring of class *i* of an individual of class *j*, and let 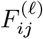 have the distribution of *F*_*ij*_ biased by *F*_*ℓj*_, that is, 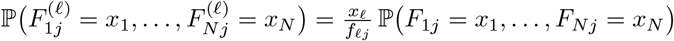. Finally, let 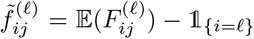 where 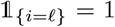 if *i* = *ℓ* and 0 otherwise). With these quantities, the expected number of class-*i*(*g, q*)-kin of *ego* through same-litter sisters of its (*g* − 1)-th ancestor is

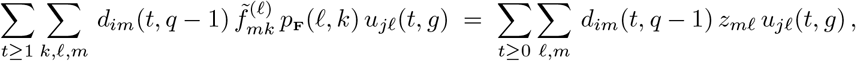

where 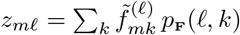. In matrix form, this is the (*i, j*)-th entry of

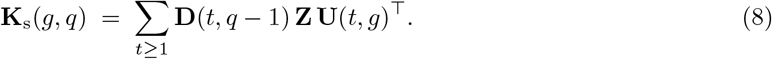

What makes Equation (8) relevant is that the matrix **Z** has a concrete interpretation: as mentioned at the beginning of this section, *z*_*ij*_ is the expected number of same-litter sisters of a focal individual in class *j* that have just been born in class *i*. Thus, the matrix **Z** can be estimated from detailed census data. Alternatively, **Z** can be computed from the matrices **S** and **F** and the covariances Cov(*F*_*ij*_, *F*_*ℓj*_). Indeed, elementary calculations give

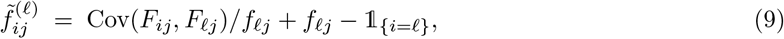

from which **Z** can then be computed. A few special cases are worth pointing out. First, when at most one offspring is produced during a projection interval, we have **Z** = **0**.

Second, if there is a single newborn class, say class 1, then *z*_*ij*_ = 0 if *i* ≠ 1 or *j* ≠ 1, and

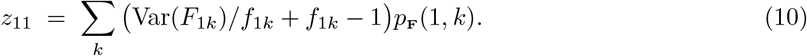

This covers the majority of the models used in practice. Third, when the numbers of offspring produced by an individual in the various classes are independent Poisson variables,

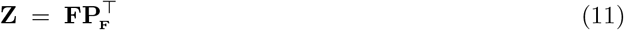

This formula can be used by default when the covariances of the fertilities are not available.

#### Expression of the kinship matrix

Combining Equations (6), (7) and (8), we get the following expression for the kinship matrix **K** = **K**_y_ + **K**_o_ + **K**_s_, which is valid for all *q* ≥ 0 but for *g* ≥ 1 only:

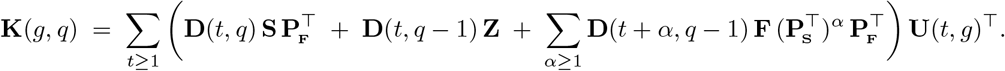

For *g* = 0, the (*g, q*)-kin of *ego* are its *q*-th descendants (and *ego* itself when *q* = 0). Thus, **K**(0, 0) = **I** and, for *q >* 0,

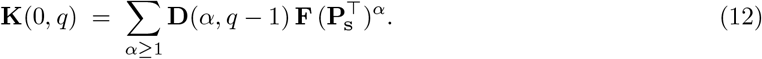

Note that, compared to Equation (7), there is no factor 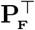 to the right of 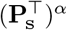.

In order to get a unique expression of **K**(*g, q*) that works for all *g ≥* 0, we let **U**(*t,* 0) = **0** for *t ≥* 1, in agreement with Equation (4) and the definition of weak compositions in Equation (3); but for *t* = 0 we use the special convention **U**(0, 0) = **I**. Setting **Y**(*g*) = **I** if *g* = 0 and 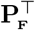 otherwise, this gives the following general expression for the kinship matrix, valid for any (*g, q*) ≠ (0, 0):

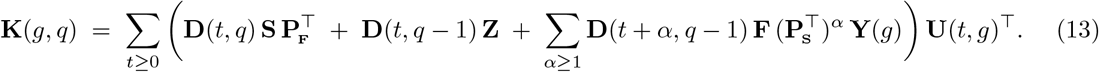

Note that for *q* = 0, this formula reduces to

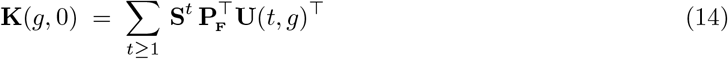

and gives the probability that the *g*-th ancestor of *ego* is alive, structured by the class of *ego* and of the *g*-th ancestor.

### 2.5 Computing the kinship matrix in practice

Equation (13) gives the kinship matrix as an infinite sum. However, in numerical applications it will have to be approximated by a finite sum, for instance of the form

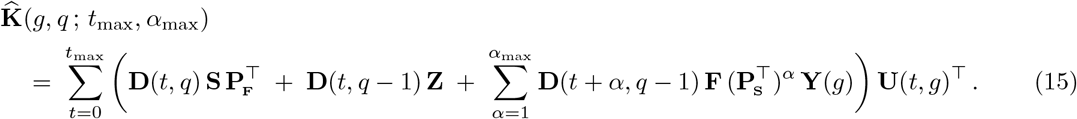

This raises the question of determining how many terms to sum, and of the precision of the resulting estimate. In SM.6, we give an explicit upper bound on 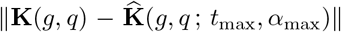 that can be used to chose *t*_max_ and *α*_max_ so as to meet a target accuracy in the estimation. A procedure to do so automatically is implemented in our code (see SM.1). It should be pointed out, however, that the upper bound of SM.6 is rather crude and that the actual precision will typically be much better than it suggests.

Importantly, when there exists a maximal age *ω* in the model (that is, *ω* such that **S**^*ω−*1^ ≠ **0** and **S**^*ω*^ = **0**), the sums in Equation (13) are, in fact, finite: indeed, in that case, **U**(*t, g*) = **0** for all *t > g ω*; **D**(*t, g*) = **0** for all *t >* (*q* + 1) *ω*; and 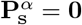 for all *α* ≥ *ω*. As a result,

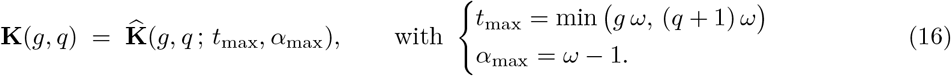

This gives a useful heuristic to choose *t*_max_ and *α*_max_ when the values obtained from the upper bound of SM.6 lead to prohibitively long calculations: first, find a reasonable estimate of the maximum age that one can expect to observe in the population, by picking 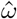 such that the probability that an individual reaches age 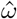 is less than some very small *ε* (e.g., by taking 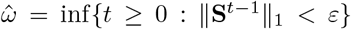}, where ||**X**||_1_ = sup_*j*_ Σ_*i*_|*x*_*ij*_|); then use the values of *t*_max_ and *α*_max_ obtained by plugging 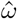 in Equation (16).

Although there is no mathematical guarantee – that is, other than that given in SM.6 – on the difference between **K** and 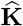 when using 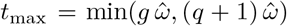 and 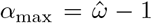, this difference should be well within the modelling error of the projection model, as it will exclusively be due to individuals that live past 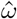 – which are allowed by the model but may not exist in practice. One way to test this is to compare the results obtained with the projection matrix **A** to those obtained with a model 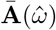 where the maximal age has been capped (as discussed in SM.7). The biological relevance of 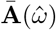 compared to that of **A** is something that can be assessed independently; and once this is done, the kinship matrix of 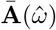 can be computed exactly using 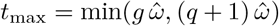 and 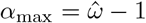.

### 2.6 Aggregated measures of kinship and relatedness

Having a general expression for the number of any type of kin makes it possible to consider various aggregated measures of the kinship structure of a population. For instance, the matrix

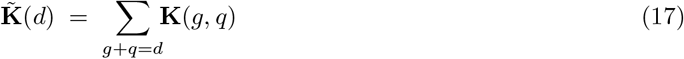

gives the number of kin that are at distance *d* from *ego* in their genealogical tree. If *λ* > 1, 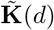 diverges as *d* → ∞, whereas if *λ* < 1 it goes to zero. It is thus natural to wonder how 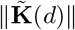 grows/decreases as a function of *d*. In SM.9, we show that it grows like 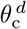, where

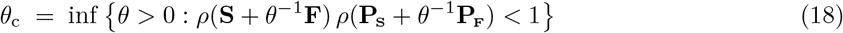

and where *ρ* denotes the spectral radius. This characterization makes it straightforward to compute *θ*_*c*_ numerically. In SM.9 we also explain why we expect to have

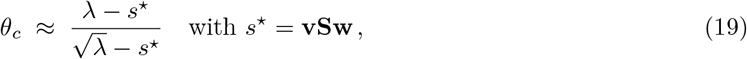

where the vector of reproductive values **v** and the stable distribution **w** are such that **vw** = 1.

One of the notable feature of *θ*_c_ is that it describes the global kinship structure, at a large-scale. This contrasts with the matrices 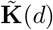, which can only be computed for relatively small values of *d* (see SM.10) and thus only give us access to the local, small-scale structure of the kinship.

In applications, *θ*_c_ could be a useful descriptor for questions ranging from life history theory to inclusive fitness: for instance, Equation (19) shows that for a fixed value of *λ* > 1, the number of kin is maximized by increasing *s*\*, i.e. by *K*-strategies; but that if *λ* < 1 it is optimized by decreasing *s*\*, i.e. by *r*-strategies. The statistic *θ*_*c*_ could thus be used to compare life history patterns in terms of the *r*–*K* continuum, as an alternative to the generation time *T* = 1/(*λ* − *s*\*) (Bienvenu and Legendre, 2015). In the context of inclusive fitness, since the matrices **K** can be interpreted as measures of relatedness (see SM.11), *θ*_*c*_ is an indicator of how much relatedness is carried by various kin: for instance, the higher *θ*_c_, the more relatedness is carried by close kin – but the smaller the relative contribution of close kin compared to that of distant ones.

Of course, 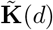 and *θ*_c_ are only two examples of aggregated measures of kinship that can be defined from the matrices **K**. In general, one may want to consider various statistics of the form Σ_*g,q*_ *f*(**K**(*g, q*), *g, q*), for functions *f* and values of (*g, q*) that depend on the specific application. If the application requires summing on large values of *g* and *q*, this may not be possible because of prohibitive computation times. In that case, one possible strategy could be to first turn the projection matrix into a 1 × 1 matrix projecting the unstructured population (see Bienvenu et al., 2017; Coste et al., 2017), and then use the efficiently computable formulas of **K**(*g, q*) for 1×1 models given in SM.8.

##### Box 2: Illustration

We illustrate our method on the most common model in animal ecology: the extended Leslie model, i.e. an age-structured model with a class that regroups all individuals older than a certain age. More complex examples are given in SM.2, to illustrate and discuss specific points; but once an appropriate projection matrix has been obtained the method is always the same. Recall that we need to consider a pre-breeding model – that is, in the case of a 3-class model, a projection matrix of the form

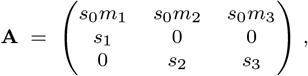

where classes 1 and 2 correspond to individuals that are about to turn 1 and 2 year(s) old, respectively, and class 3 corresponds to older individuals; *s*0 is the probability that newborns survive their first year, *s*_*i*_ the probability that class-*i* individuals survive to the next year and *m*_*i*_ the average number of (female) newborns to which they give birth.

Using the life table for the ground squirrel *Spermophilus dauricus* from Luo and Fox (1990), we get the following survival and fertility matrices:

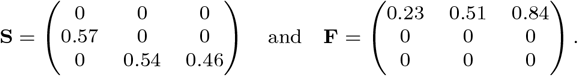

The asymptotic growth rate is *λ* = 1.00 and the stable class distribution is **w** = (0.47, 0.27, 0.27). From this, Equation (2) gives

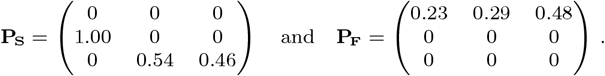

For instance, the (3,2)-th entry of **P**_**S**_ shows that, at the demographic steady state, 54% of class-3 individuals were in class 2 the previous year. Similarly, the (1, 1)-th entry of **P**_**F**_ indicates that a yearling has a 23% probability of having been born to a yearling the previous year. Since there is a single newborn class, we can use Equation (10) to compute the matrix **Z** under arbitrary assumptions on the fertilities. But here, in the absence of additional information, we will simply assume that they correspond to Poisson random variables, so that 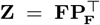. This yields

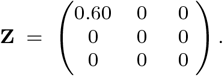

Let us now compute **K**(1, 0), whose (*i, j*)-th entry is the probability that the mother of an individual of class *j* is currently alive and in class *i*. Using *t*_max_ = 9 and *α*_max_ = 0, which as per SM.6 gives an approximation error 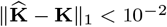, we get

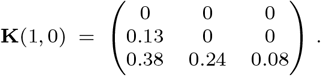

This matrix indicates, for instance, that a yearling has a 13% probability that its mother is alive in class 2, and 38% that it is alive in class 3. Note that kin are structured by their *current* class. Thus, it is not possible to have yearling mothers here: even if the mother of *ego* was a yearling when *ego* was born, by now it will have moved to an adult class.

Figure 3a gives a graphical representation of **K**(2, 1), the kinship matrix for aunts. The total number of aunts (irrespective of their class) of an individual sampled according to the stable class distribution is **1K**(2, 1)**w** = 0.36, and numerically it is possible to compute its elasticities to the entries of the projection matrix, i.e. *∂* ln **1K**(2, 1)**w***/∂* ln *a*_*ij*_. These are plotted in Figure 3b, which shows that the number of aunts is most affected by relative changes in the survival probabilities *s*_1_ and *s*_3_.

**Figure 3:**
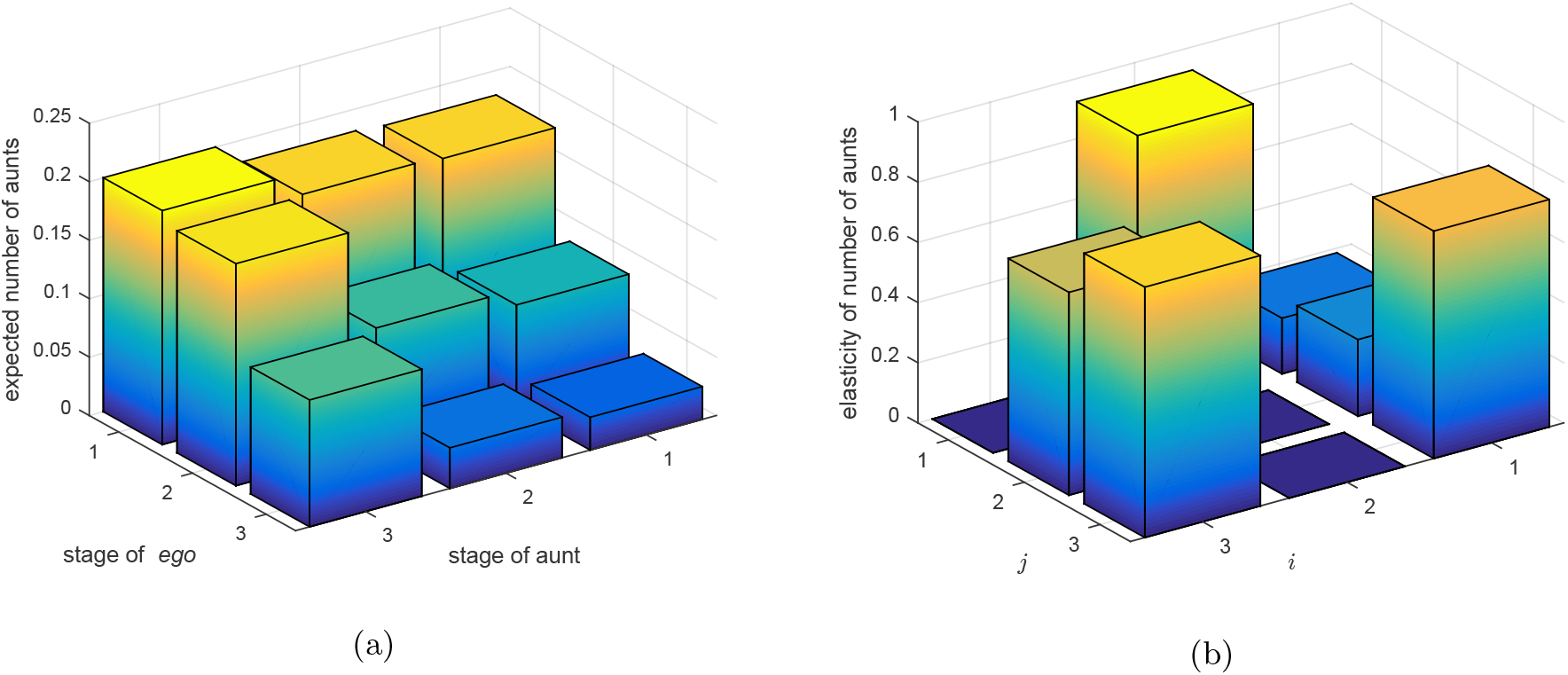
(a) graphical representation of the matrix **K**(2, 1), whose entries give the expected number of aunts of *ego* as a function of its class and that of its aunts; (b) the elasticities of the expected total number of aunts of *ego* (sampled at random in the stable population), *∂* ln **1K**(2, 1)**w***/∂* ln *a*_*ij*_.

Finally, we get sensibly the same results when taking the values of *t*_max_ and *α*_max_ suggested by Equation (16), with the maximal age 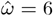 reported in Luo and Fox (1990). This is also the case with other models that we studied (see SM.2), which tends to suggest our simple heuristic to chose *t*_max_ and *α*_max_ based on a maximal age is adequate.

In SM.12, we compare the kinship matrices **K**(1, 0) and **K**(2, 1) to the output of individual-based simulations, confirming our formulas.

## 3 Discussion

In this article, we have developed a unified framework to compute the expected number of kin of a focal individual (as a function of the vital traits of the focal individual and of its kin), directly from the demographic rates of a population. The main feature of our method is that it applies to any kin relationship (mother-daughter, aunt-niece, cousin-cousin, etc) and any structured population described by a matrix population model – be it structured by age (as illustrated in Box 2), by stage (SM.2.1), by patch (SM.2.3), or by any conceivable type of discrete class. These theoretical results come with readily usable implementations that make them available to any ecologist. Here we discuss the main limitations and implications of this unified framework, and give some perspectives for future research.

### Conceptual and practical limitations

Our work is set in the framework of matrix population models, and thus inherits its usual limitations. In particular, density-dependence and environmental stochasticity – which are not considered by our input model – are likely to be two of the main sources of discrepancies between our theoretical predictions and reality. However, because our mathematical reasoning is based directly on the stochastic model underlying the matrix population model (rather than on its deterministic description of the average population dynamics), demographic stochasticity is not a limitation: it is taken into account in our calculations, and although our final formulas correspond to expected values with respect to this stochasticity, our methods could also be used to compute, e.g, variances. This is discussed below.

Another important limitation of our approach is that it hinges on the assumption that the dynamics of the population has been at the steady state for an extended period of time. This limits the validity of our formulas in populations whose demography has been perturbed in the recent past – all the more so when studying distant kin relationships. Unfortunately, there does not seem to be an easy way to circumvent this using our techniques, and there may not be alternatives to recursive formulas for such models (Caswell and Song, 2021).

When applying our results, one should also pay attention to the fact that they are formulated for models where the fertilities are independent from adult survival. This assumption is required for the connection with the stochastic population model that implicitly underlies our calculations, but it is not compatible with post-breeding models. However, these can (usually easily) be reformulated as pre-breeding models, so this is not a major limitation of our work.

As implicit with matrix population models, we work in the context of monoparental genealogies. This setting could correspond either to a one-sex species, or to a two-sex species with a female-based demographic model. In the latter case, it should be noted that our formulas are based on a notion of kinship that only considers matrilines, and will thus differ from the traditional notion of kinship in a complete pedigree: for instance, the daughters of the brothers of the focal individual will not be counted as its nieces. Although this is not a limitation of our results, this is something important to have in mind when interpreting them. Incidentally, this specific notion of kinship can in some contexts be interpreted as a direct measure of relatedness. Indeed, as explained in SM.11, in an idealized setting (namely, a large panmictic population where males and females have similar life-histories and where no two individuals have the same two parents), discarding kin that are not based on matrilines has the same effect as the dilution of genetic material over successive generations. Thus, although our results were first and foremost developed to study kinship, they could have wider-ranging implications in evolutionary biology, by working as a measure of relatedness.

Finally, a practical limitation of our results is that the computing time of our formulas increases rapidly when we consider increasingly distant kin, and can become prohibitively long – especially for organisms with a “slow” strategy on the slow-fast continuum. Because close kin are what matter in most situations, this is, in general, hardly going to be a problem. However, should very distant kin be of interest, we provide two partial solutions to prohibitive computing times: first, in Section 2.6 we use our results to derive an efficiently computable statistic to describe the large-scale kinship structure of a population; second, if one is ready to discard the structure of the population, then in SM.8 we give a simple formula for the expected number of kin that is trivial to compute in practice. Note that both of these results are made possible by the fact that we have a generic formula for the kinship, and would presumably not have been possible using ad hoc formulas for each type of kin.

### Implications and perspectives

First and foremost, having a generic formula for arbitrary kin relationship makes it possible – and elementary – to define tailor-made measures of kinship to be used in a specific setting. For instance, anthropologists who would like to compare the number of lineal vs collateral kin can easily do so by summing the kinship matrices of their choice – or, more generally, combine them however they want to produce the best suited metric for their particular application.

Our methods can be used to compute, at least numerically, the sensitivity or the elasticity of the kinship structure to the traits implemented in the model. This is illustrated on a concrete example in Box 2. Because our formulas apply to any primitive projection matrix, they can be used to study models where classes are based on several traits, such as metapopulation models (Lebreton and Gonzáles-Dávila, 1993) or more general multitrait models (Roth and Caswell, 2016; Coste et al., 2017; Coste and Pavard, 2020). This makes it possible to study and compare the relative effects of different traits on the kinship. We illustrate this in SM.2.3, where we use a *stage × patch* model to study the effect of dispersal.

Finally, because our results make new use of classic tools to solve a long-standing problem, they could spur further theoretical developments in the field of matrix population models. An example of this is given by our introduction, in SM.5, of the previous-generation matrix (which is a dual version of the classic next-generation matrix of Cushing and Zhou, 1994) and of the Euler–Lotka matrices (which give two simple generalizations of the Euler–Lotka equation to arbitrarily structured populations). Despite being very natural from a mathematical and biological point of view, these quantities do not seem to have been identified as such before (although similar ideas can be found in Lebreton, 2005). Though not directly linked to kinship, these results are straightforward consequences of the framework we have developed to study it.

Our work offers several perspectives in terms of extensions, in particular to see if our methods can be extended beyond the framework of matrix population models. A natural candidate for this are integral projection models (Rees et al., 2014). Because the variations in the individual realizations of the vital rates have a relevant impact on kin structure (Coresh and Goldman, 1988; Tuljapurkar et al., 2020, 2021), another perspective is to go the beyond the “expected-value” description of the kinship structure given here – namely by deriving statistics that describe the demographic stochasticity underlying our calculations, instead of averaging it out. For instance, with our method, the classic results of Everett and Ulam (1948) on multitype Galton–Watson processes could be used to compute the variance of the number of kin. Finally, an important line of research that we plan to explore is the extension of our work to two-sex models. This is particularly challenging, as it requires leaving the framework of monoparental genealogies and will therefore have to be done using a combination of analytical and numerical methods.

## Acknowledgments

This study is supported by a grant from the Agence Nationale de la Recherche (ANR-18-CE02-0011, MathKinD). CFDC was funded by Centre of Excellence grant from the Research Council of Norway (SFF-III, project no. 223257). FB was partly funded by Agence Nationale de la Recherche (ANR-16-CE27-0013), EPSRC Fellowship EP/N004833/1 and a Transilvania Fellowship from the Transilvania University of Bras, ov. We thank Arpat Ozgul for sharing data on the life cycle of the yellow-bellied marmot (Marmota flaviventris).

## Supplementary Materials

### SM.1 Python and MATLAB implementations of the formulas

We provide two implementations of the methods presented in this paper:

- A MATLAB implementation that can be used to reproduce all numerical calculations and figures presented in this manuscript. Using it requires knowing MATLAB.
- A Python implementation with a flexible, general-purpose interface. Python is a free, open-source programming language that is available on every platform. This implementation provides a well-documented library that can be used in Python scripts, but it also comes with a graphical user interface that requires no prior knowledge of any programming language to be used (see Figure 4).

**Figure 4:**
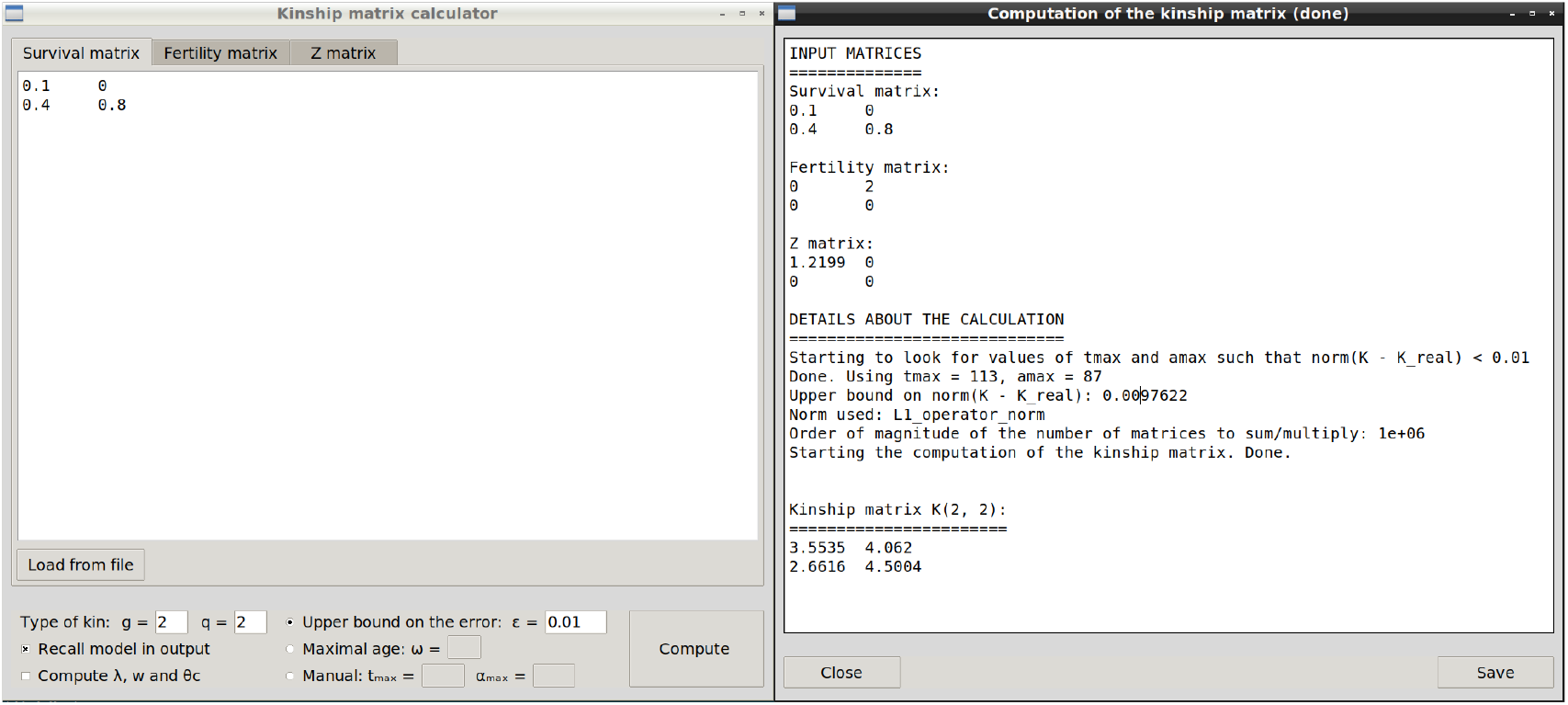
Graphical user interface provided with the Python implementation of the formulas.

Both implementations can be downloaded with detailed instructions from 10.5281/zenodo.4680715.

### SM.2 Additional illustrations

In Box 2, we have illustrated our method using an extended Leslie model. This choice is natural, because these are the most widely used models in animal ecology. However, this does not illustrate the fact that our method works for any population structure. This also did not give us the occasion to address relevant questions that can arise in some settings. Thus, in this section we give the following additional illustrations:

- The classic life cycle of the killer whale (*Orcinus orca*) from Brault and Caswell (1993). This is a fairly extreme example of how a “slow” strategy (high survival, low reproduction) can make the calculations computationally intensive. It is also used to discuss other questions (birth-flow and post-reproductive classes).
- The complex life cycle of the yellow-bellied marmot (*Marmota flaviventris*) from Ozgul et al. (2009). This is an example of a post-breeding model whose conversion to a pre-breeding model is non-trivial.
- A generic metapopulation model that illustrates how our methods can be used to study the effects of traits such as dispersal on kinship.

#### SM.2.1 A slow strategy: the killer whale *Orcinus orca*

Consider the following classic projection matrix for the killer whale *Orcinus orca*, taken from Brault and Caswell (1993):

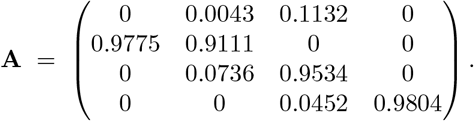

This is a so-called Lefkovitch matrix with four stages (newborn, juvenile, reproductive adult and post-reproductive adult).

##### Birth-flow

This model is what is known as a *birth-flow* model: since orcas do not have a fixed breeding season, individuals can be born at any time of the year. As a result of this, the fertilities (first row of the projection matrix) incorporate a part of the survival of the mother; and thus the number of offspring that an individual produces between time *t* and *t* + 1 *cannot* rigorously be independent from its survival during that period.

However, one should keep in mind that, even in pre-breeding models, the independence of survival and reproduction is never perfectly met in practice because there is always a trade-off between survival and reproduction. Thus, just as any of the many working hypotheses of matrix population models, this independence is an “ideal” mathematical assumption that is required for the calculations to be exact and whose relevance will depend on the biological situation at hand. In a post-breeding model, it is typically not acceptable (especially given that it is almost always possible to convert such a model into a more suitable pre-breeding one); but in a birth-flow model, it can be – all the more so when survival is high and fertility is low, which tends to decorrelate them. The intuition for this is the following: when survival is high and reproduction is low, the death of pregnant mothers (which is the source of the non-independence that is problematic to us) is rare and can be neglected.

As a more formal argument, let *S*_*i*_ be the Bernoulli variable corresponding to the survival of an individual from class *i* from one year to the next (thus, letting *p*_*i*_ = ℙ(*S*_*i*_ = 1), here *p*_2_ = 0.9847 and *p*_3_ = 0.9986), and let *F*_*i*_ be the number of offspring it leaves in the next year. Since orcas produce at most one offspring per year, *F*_*i*_ is a Bernoulli variable with parameter *f*_*i*_ (here with *f*_2_ = 0.0043 and *f*_3_ = 0.1132). Assuming that birth and death occur at constant rate (with at most one birth), standard calculations yield that 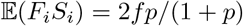. Thus, the correlation coefficient between *S*_*i*_ and *F*_*i*_ is

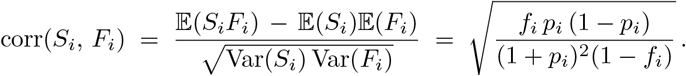

Numerically, this gives corr(*S*_2_, *F*_2_) = 0.0041 and corr(*S*_3_*, F*_3_) = 0.0067, which tends to confirm our intuition about the low dependence between *S*_*i*_ and *F*_*i*_. As a result, for our purposes the pre-breeding model gives a decent approximation of the birth-flow one and can be used to compute **K**.

##### Reducibility

Here, the projection matrix **A** is *not* primitive, because of the post-reproductive class. However, it still has a dominant eigenvalue *λ* = 1.0254 with a unique associated right-eigenvector, namely **w** = (0.0370, 0.3161, 0.3229, 0.3240). Since **w** > 0, as explained in SM.3 it is still possible to use Equation (2) to compute the genealogical matrices, yielding

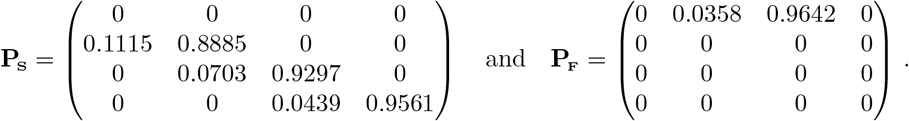

##### Estimation of the kinship matrices

First, we need to compute the same-litter newborn sisters matrix **Z** (see Section 2.4). Since orcas give birth to at most one offspring a year, **Z** = **0**.

Next, we need to choose appropriate values of *t*_max_ and *α*_max_. Here however, because of the high survival and low reproduction, the values of (*t*_max_*, α*_max_) suggested by the upper bound of SM.6 are not practical: indeed, for the *ℓ*_1_ operator norm 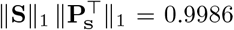, so that we would need to take, e.g., *t*_max_ = 9630 and *α*_max_ = 9523 to compute 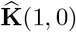 with the guarantee that 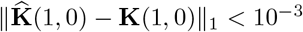. This leads to prohibitively long calculations. For most other classic norms, 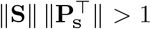 so our upper bound on the error does not even converge.

Therefore, we use 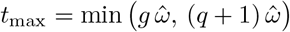 and 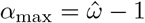, as suggested in Section 2.5, with the maximum age 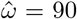 reported by Brault and Caswell (1993). With this, we get the following matrices for mothers and grand-mothers, respectively

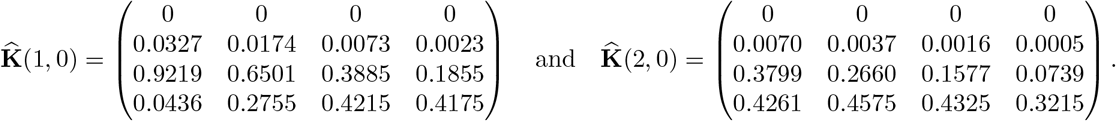

For aunts and cousins, we get

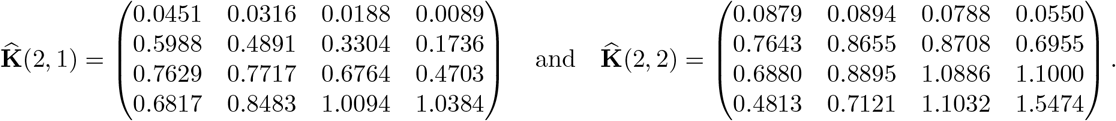

In the absence of useful upper bound on the error 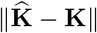, to get an idea of the reliability of these estimates we can proceed as suggested in Section 2.5 and compare them with the matrices obtained from the “capped” *age stage* model **Ā**(90) described in SM.7. Indeed, in the capped model the corresponding kinship matrices 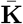 can be computed exactly using finite sums.

Here, the capped matrix **Ā**(90) is a 265 × 265 matrix. Whether it gives an appropriate description of the population can be assessed by comparing its descriptors to those of **A** (for instance, 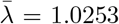 vs *λ* = 1.0254 or 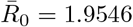 vs *R*_0_ = 2.0131, etc). One can also “fold” it over age to recover a model with the same population structure as **A** (see Bienvenu et al., 2017; Coste et al., 2017). This gives

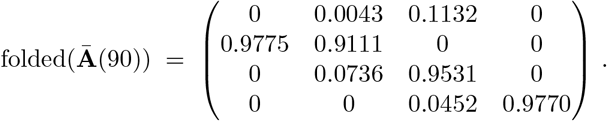

As expected, the survivals are slightly lower than in **A**, in particular for post-reproductive adults (0.9770 vs 0.9804). But, overall, the comparison suggests that here the initial model **A** and the capped model **Ā**(90) describe fairly similar population processes.

Comparing the kinship matrices 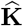 corresponding to **A** with the kinship matrices 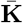 corresponding to **Ā**(90) shows that neglecting individuals over 90 year old has a slight but acceptable impact on the estimation; compare for instance the total number of mothers in **A** vs in **Ā**(90), i.e. 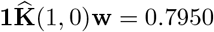 vs 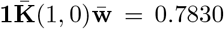 of grand-mothers (0.5793 vs 0.5232); of aunts (1.9590 vs 1.9005); of cousins (2.9983 vs 2.9390), etc.

##### Comparison with another cetacean

The kinship matrices can be used to quantitatively compare the familial structure of orcas with that of other species. For instance, for the North Atlantic right whale (*Eubalaena glacialis*), using the projection matrix from Young and Keith (2011),

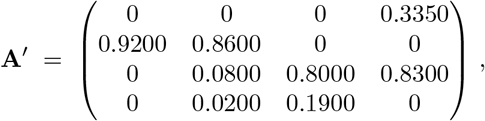

we get the following kinship matrices for aunts and cousins:

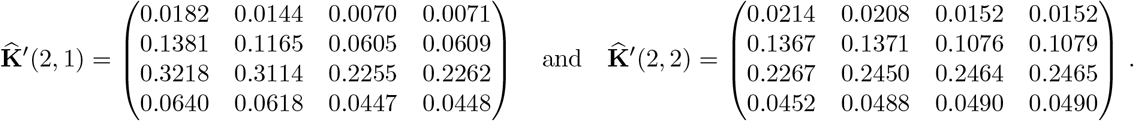

Despite similar growth rates for the two populations (*λ* = 1.0254 vs *λ′* = 1.0000), we see that killer whales have far more kin than North Atlantic right whales: compare for instance the total number of aunts 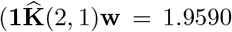 vs 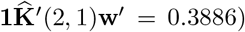 or of cousins (2.9983 vs 0.4273) of *ego* in both models. This general trend is confirmed by the fact that *θ*_c_ = 1.3778 and *θ′*_c_ = 0.9996 (cf Section 2.6).

This is unsurprising and can be related to the very different social structures of both species: indeed, killer whales live in pods with their kin and hunt cooperatively with them, whereas North Atlantic right whales are more solitary. Thus, there likely is a greater selective pressure for killer whales to have more kin than for North Atlantic right whales.

#### SM.2.2 A complex life cycle: the yellow-bellied marmot *Marmota flaviventris*

Consider the following life cycle for the marmot *Marmota flaviventris* from Ozgul et al. (2009):

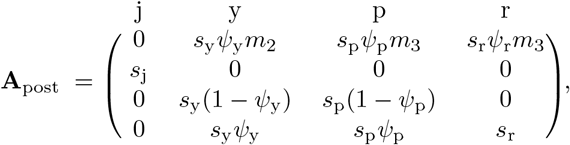

where the four classes are juveniles, yearlings, pre-reproductive adults and reproductive adults; *s*_*x*_ is the probability that an individual of class *x* survives the year, *ψ*_*x*_ the probability that it reproduces at the end of it, and *m*_2_ (resp. *m*_3_) litter size of a 2 year-old (resp. ≥3 year-old). This models is a good example of the non-conventional (that is, neither Leslie nor strictly Lefkovitch) models that are increasingly used in animal ecology. As the subscript suggests, it is a post-breeding model. The non-independence between reproduction and survival is made apparent by the fact that factors such as *s*_*x*_ or *ψ*_*x*_ in the fertility of class *x* also appear in the third and/or fourth row of column *x*.

The reason why we present this model is that the conversion to a pre-breeding model is non-trivial and may at first glance seem to be an obstacle to the calculation the kinship matrices with our method. However, this is not the case: all we need to do is split the “reproductive adults” class into three subclasses so as to distinguish between reproductive adults based on whether they were yearlings, pre-reproductive adults or reproductive adults the previous year. Once this is done, the corresponding pre-breeding model is easily seen to be

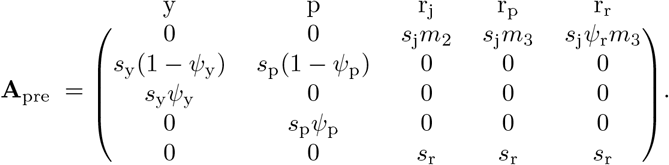

Here the classes are yearling, pre-reproductive adults, reproductive adults that were yearling the previous year, reproductive adults that were pre-reproductive adults the previous year, and reproductive adults that were already reproductive adults the previous year.

Once the projection matrix **A**_pre_ has been obtained, the kinship matrices can be computed exactly as in Box 2 or in SM.2, without regard to the unusual form of the life cycle. Using the numerical values from Ozgul et al. (2009), we have the survival and fertility matrices

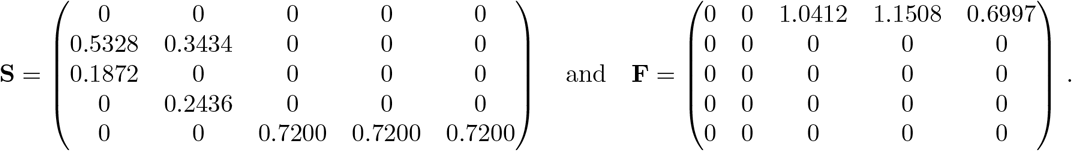

Assuming Poisson reproduction, we get – for instance – the following kinship matrix for sisters:

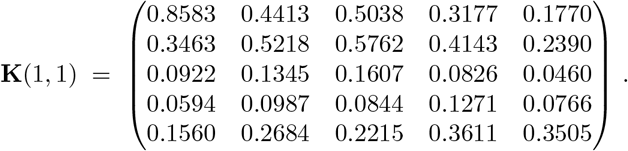

If we want a coarser description of the population and would like to get the number of kin structured by the original classes of the models, if suffices to aggregate the classes r_y_, r_p_ and r_r_, i.e. to sum the corresponding entries after weighting each column by its asymptotic abundance. This yields:

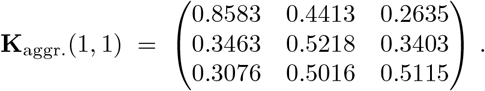

Note that it is not possible to recover the number of kin of juveniles nor the number of juvenile kin, because these are not counted by the model.

#### SM.2.3 A metapopulation model: studying the impact of dispersal on kinship

Since our formulas apply to a very large class of projection models (see SM.3), they can be used to study models that are based on combinations of traits (referred to as multitrait models in Coste et al. (2017) and hyperstate models in Roth and Caswell (2016)), in particular metapopulation models (sensu Lebreton, 1990).

##### The model

Let us consider the following metapopulation model with two stages (juveniles and adults) and two patches (A and B), where juveniles can disperse:

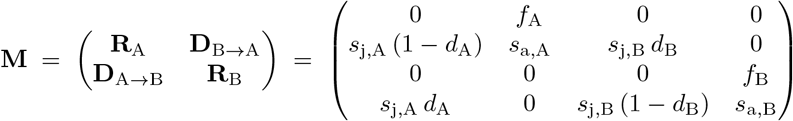

For such a model, the kinship matrices can be computed from **M** in order to understand the kinship structure within and across patches. To understand the effect of the metapopulation structure, we can compare it to the kinship structure associated to each patch in the absence of dispersal.

In order to work out a numerical example, let us fix some simple parameters for patches A and B. We choose them to correspond to a situation where patch A has slightly better probabilities of survival and of reproduction (say, *s*_j,A_ = 0.4, *s*_a,A_ = 0.5, *d*_A_ = 0.5 and *f*_A_ = 0.8 × 2.3 for patch A, where *f*_A_ is the product of the probability of reproduction and of the average number of offspring that survive their first year, conditional on reproduction; and *s*_j,B_ = 0.3, *s*_a,B_ = 0.4, *d*_B_ = 0.4 and *f*_B_ = 0.7 × 2.3 for patch B). This gives the projection matrix

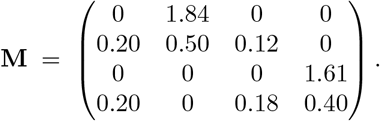

The corresponding genealogical matrices are

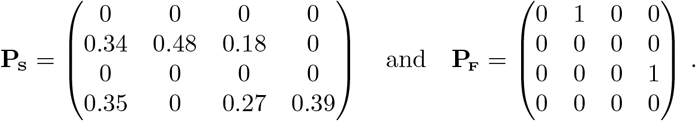

Finally, assuming Poisson reproduction, Equation (11) from 2.4 gives the following same-litter newborn sisters matrix:

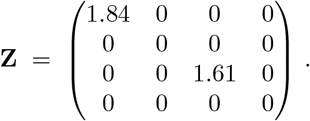

##### The previous generation matrix

Because there are several newborn classes here, the previous-generation matrix **G** (see SM.5) has an interesting structure:

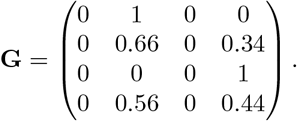

This shows that adults found in patch A are likely have been born in that patch (66% probability), whereas in patch B they are more likely to have been born the other patch and dispersed from it (56% probability). If we look at *g*-th generation ancestors, then

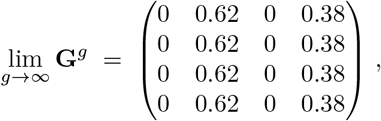

meaning that a distant ancestor of *ego* has a 62% probability of having been born in patch A, irrespective of the stage or the patch of *ego*; in fact, the ancestral lineage of *ego* goes to and fro between patches A and B, spending on average 62% of its time in patch A.

##### Quantifying the effect of dispersal on kinship

Let **K**_ref_ be the kinship matrix in the absence of dispersal (i.e. *d*_A_ = *d*_B_ = 0) and let Δ**K** = **K − K**_ref_. The matrix Δ**K** enables us to quantify the effect of dispersal on the kinship structure. For instance, looking at cousins,

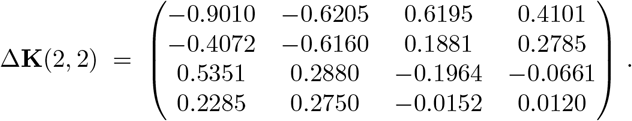

As expected, dispersal not only decreases the number of cousins within patch A, but it also decreases the overall number of cousins for individuals from patch A: if for *x* ∈ {j, a} and *X* ∈ {A, B} we let Δ_tot_(*x, X*) = Σ_(*y,Y*)_ Δ*k*_(*y,Y*),(*x,X*)_ denote the total number of kin (of any age and of any patch) that an individual in class *x* and patch *X* gains as a result of dispersion (where Δ*k*_(*y,Y*),(*x,X*)_ denotes the entries of Δ**K**); in other words, Δ_tot_(*x, X*) is the sum of the elements in the (*x, X*) column of Δ**K**), then we see that Δ_tot_(j, A) = −0.5447 < 0 and Δ_tot_(a, A) = −0.6735 *<* 0. By contrast, although dispersal decreases the number of within-patch cousins for patch B, it actually increases the overall number of cousins for individuals from that patch: Δ_tot_(j, B) = 0.5960 *>* 0 and Δ_tot_(a, B) = 0.6346 *>* 0.

If instead of focusing on a particular kin relation (here cousins) we want to have a more global vision of this, we can compare *θ*_c_(**M**) = 1.0544, 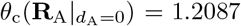 and 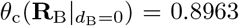. Again this shows that dispersal tends to decrease the number of kin for patch A and for the overall population, but that it increases it for patch B. Note that these values have to be interpreted in light of the growth rates *λ*(**M**) = 1.0382, 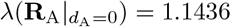 and 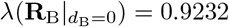.

Finally, it is also possible to compute the sensitivities of the kinship structure to dispersal. For instance,

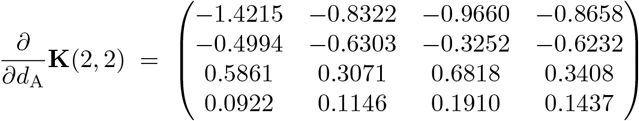

and

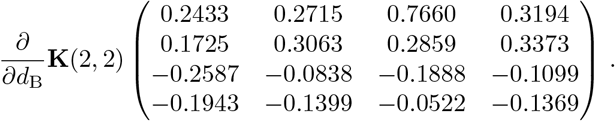

Similarly, *∂θ*_c_*/∂d*_A_ = −0.2163 and *∂θ*_c_*/∂d*_B_ = 0.1384.

In summary, this simple toy model shows that our methods give a precise and flexible way to quantify the effect of various life-history traits such as dispersal on kin structure.

##### Effect of categories on kinship

Another way to study the effect of various traits on kinship is to compare a model that implements a trait with the corresponding model that does not implement it (the “folded” model, see Coste et al., 2017; Coste and Pavard, 2020). Here, the model is based on two traits: stage and patch. Patch-related traits account for dispersal and the heterogeneity in vital rates between patches, whereas stage-related traits account for the heterogeneity in vital rates between stages. We assess the overall effect of each of these traits on kinship by comparing the kinship matrices of various models: the original metapopulation model **A** implementing boths traits; the folded-over-patch model; the folded-over-stage model; and the unstructured model (i.e. the corresponding 1×1 matrix). The corresponding values of the expected number of aunts and cousins, as well as the probability that the mother is alive, are given in Table 1. They show that adding the patch structure to the unstructured model or to the stage-based model has little effect on kin frequencies. By contrast, adding the stage structure to the the unstructured model or to the patch-based model has a noticeable effect. This suggests that there is more heterogeneity between stages than between patches, and that stage is a more relevant trait for the kinship structure than patch.

**Table 1:**
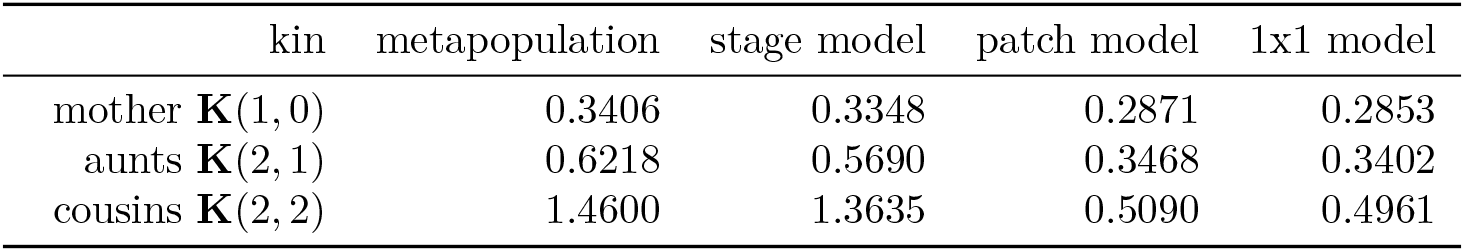
Probability that the mother of *ego* is alive and expected number of aunts / cousins of *ego*, for four models: the stage×patch metapopulation, the corresponding stage-based model, the corresponding patch-based model, and the corresponding unstructured model.

### SM.3 Reducible projection matrices

In order for the genealogical matrices **P**, **P**_**S**_ and **P**_**F**_ given by Equations (1) and (2) to be well-defined, the projection matrix **A** must have:

i. a positive eigenvalue *λ* that is greater in modulus than any other eigenvalue;
ii. a unique right-eigenvector **w** associated with *λ*;
iii. **w***>* 0.

If **A** is primitive – that is, irreductible and aperiodic – then the Perron-Frobenius theorem ensures that this is the case (see e.g. Meyer, 2000; Varga, 2000). Most population projection matrices encountered in practice are primitive, and thus we have relied on primivity to state our results in the main text, for simplificity. However, there are biologically relevant situations in which a non-primitive matrix may verify points (i–iii). Examples of such situations include, among others, the presence of post-reproductive classes (as is the case for the life-cycle of the killer whale *Orcinus orca* presented in SM.2.1); and meta-population models with “sink” populations.

In practice, the easiest way to determine whether our results can be applied is to compute the eigen-elements of the projection matrix and see whether points (i–iii) are verified. If points (i) and (ii) are verified but the dominant eigenvector **w** has some zero entries, then it is possible to remove the corresponding classes from the model and proceed with the resulting projection matrix.

In the rest of this section, we recall some elementary properties of reducible matrices that are helpful to easily identify classes of matrices for which the genealogical Markov chain is defined – and to better understand its ergodicity (or lack thereof). These results will also be useful in our study of the Euler– Lotka matrices in SM.5 and of the “capped-age” model of SM.7.

Assume that we have a reducible projection matrix **A**. In that case, the life cycle can be decomposed into several irreducible components, with “flows” of individuals between these components. Formally, the projection matrix **A** can be written as a block triangular matrix by permuting its rows and columns:

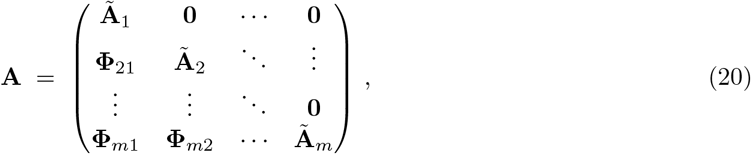

yielding what is sometimes known as a *normal form* of the matrix (see e.g. Varga, 2000, Section 2.3). In this normal form, the square matrices **Ã**_1_, …, **Ã**_*m*_ correspond to the internal dynamics of the irreducible components of the life-cycle graph, and the matrices Φ_*kℓ*_ to the flows between those components. The eigenvalues of **A** are the eigenvalues of **Ã**_1_, …, **Ã**_*m*_.

Assume that one of the matrices **Ã**_1_, …, **Ã**_*m*_, say **Ã**_*x*_, is aperiodic – and therefore primitive – and that its dominant eigenvalue is greater in modulus than any of the eigenvalues of the other matrices **Ã**_*k*_, *k* ≠ *x*. In that case, the internal dynamics of **Ã**_*x*_ will exponentially dominate that of the other matrices **Ã**_*k*_. Components that do not receive individuals from **Ã**_*x*_ will not contribute to the asymptotic constitution of the population; components that do will – but without affecting the exponential growth rate of the population. Note that the matrices **Ã**_*k*_, *k* ≠ *x*, do not need to be aperiodic: the aperiodicity of **Ã**_*x*_ will “kill” any possible periodic behavior in the components that receive individuals from it (and, while the components that do not receive individuals from **Ã**_*x*_ may exhibit a periodic behavior, they do not contribute to the asymptotic composition of the population).

Formally, let *λ* be the dominant eigenvalue of the primitive matrix **Ã**_*x*_, with *λ > ρ*(**Ã**_*x*_) for all *k* ≠ *x*, and let 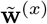 be the associated right-eigenvector of **Ã**_*x*_. Then, *λ* is an eigenvalue of **A** that is larger in modulus than any of its other eigenvalues, and the only associated (unscaled) right-eigenvector of **A** is **w** = (*w*_*i*_), with

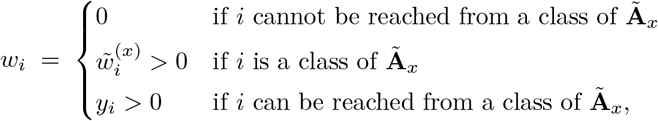

where by “*i* can be reached from *j*” we mean that there exists a a directed path from *j* to *i* in the life-cycle graph that corresponds to **A**.

A prototypical example of this situation is given by the matrix

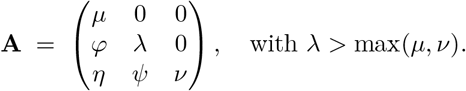

Here, the irreducible components are the 1×1 matrices **Ã**_1_ = (*µ*), **Ã**_2_ = (*λ*) and **Ã**_3_ = (*ν*). The component with the largest eigenvalue is **Ã**_2_, and it is (trivially) primitive. **Ã**_1_ sends individuals to **Ã**_2_ through *ϕ* and to **Ã**_3_ through *ν*; and **Ã**_2_ sends individuals to **Ã**_3_ through *ψ*. It is easy to check that the right-eigenvector of **A** associated with *λ* is

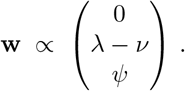

As expected, *w*_1_ = 0 because **Ã**_1_ does not receive individuals from the dominant component **Ã**_2_, and *w*_3_ > 0 if and only if *ψ* > 0, i.e. if and only if **Ã**_3_ receives individuals from **Ã**_2_. Note that neither the internal dynamics of **Ã**_1_ (that is, *μ*) nor the flows of individuals from it (that is, *ϕ* and *η*) appear in the expression of **w**.

Let us close this section by recapitulating what to do in case the projection matrix is not primitive:

1. Identify the irreducible components and find the *dominant component*, i.e. the irreducible component with the largest eigenvalue. If there are several such irreducible components, our results do not apply because the stationary distribution (if it exists) will depend on the initial composition of the population.
2. If the dominant component is periodic, our results do not apply because there is no stationary distribution.
3. If the dominant component is aperiodic, then separate the projection matrix into two matrices:

- The matrix **B** containing all of the classes that *can* be reached from the dominant component. This matrix has the same dominant eigenvalue as the original projection matrix **A** and a unique, positive dominant eigenvector. Thus, its genealogical matrices are given by Equations (1) and (2).
- The matrix **C** containing all of the classes that *cannot* be reached from the dominant component. This matrix describes a population process taking place on a smaller scale as that of **A** and **B**. If it is not primitive, go to step 1 to study it.

The matrices **B** and **C** can then be studied independently.

### SM.4 Generating weak compositions

Our formula for the kinship matrix requires summing on weak compositions, i.e. on the sets

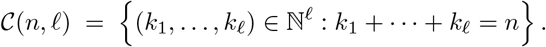

In this section, we explain how to do so efficiently.

First, note that the weak compositions of the integer *n* into *ℓ* parts are essentially the combinations of *I*_*n*+*ℓ*−1_ = {1, …, *n* + *ℓ* − 1} of size *ℓ* − 1 (that is, subsets of *I*_*n*+*ℓ−*1_ with *ℓ* − 1 elements). Indeed, the combination *C* = {*i*_1_*, ..., i*_*ℓ* − 1_} ⊂ *I*_*n*+*ℓ* − 1_, with *i*_1_ < … < *i*_*ℓ* − 1_, can be seen as the vector **u** = (*u*_*p*_) of symbols “•” and “|”, of length *n* + *ℓ* − 1, defined by: *u*_*p*_ = | if *p ∈ C* and *u*_*p*_ = • otherwise. This vector **u** can in turn be interpreted as a weak composition **k** = (*k*_1_, …, *k*_*ℓ*_) as follows:

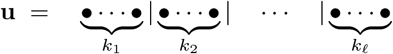

where *k*_*p*_ is the number of “•” between the (*p* − 1)-th and the *p*-th “|”. This way, *k*_1_, …, *k*_*ℓ*_ are indeed non-negative integers such that *k*_1_ + + *k*_*ℓ*_ = *n*. This shows that the number of weak compositions of *n* into *ℓ* parts is

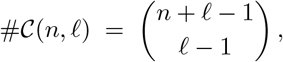

a fact that we will be of constant use in various part of this document (SM.6, SM.8, SM.9 and SM.10). More to the point here, this shows that one can obtain a weak composition **k** = (*k*_1_*, ..., k*_ℓ_) from the combination *C* = {*i*_1_, …, *i*_*ℓ*−1_}, *i*_1_ < · · · < *i*_*ℓ*−1_ by setting *i*_0_ = 0, *i*_*ℓ*_ = *n* + *ℓ*, and then

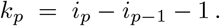

Since many programming languages – such as Python, R or MATLAB – provide a way to generate combinations, this gives a simple way to generate weak compositions.

Alternatively, the efficient Algorithm 1 below (from Knuth, 2011, Section 7.2.1.3) can be used. This approach has the advantage that it is not necessary to store the weak compositions in memory (although we do it here with the variable *list*, for the sake of giving a language-independent presentation): they can be generated one after the other as we need them and discarded as we go (typically, using a *generator* in Python).

Finally, note that although the matrices 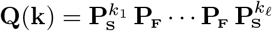 and 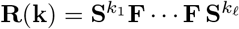 could be implemented directly from combinations rather than from compositions (by setting **Q** = **I** and then doing, for *p* = 1 to *n* + *ℓ* − 1: if *p ∈ C*, then **Q ← QP**_**F**_; else **Q ← QP**_**S**_), this approach does not take advantage of possible optimizations in the computation of matrix powers. For instance, assuming that matrix multiplication has a time complexity Θ(1) and that matrix powers are computed via exponentiation by squaring, calculating **Q** from combinations has a time complexity Θ(*n* + *ℓ*) whereas calculating it from weak compositions has a time complexity Θ(*ℓ* log(max_*p*_ *k*_*p*_)). This can make a significant difference when *n* is large and *ℓ* is small – as is typically the case in our applications.

#### Algorithm 1: Generating weak compositions

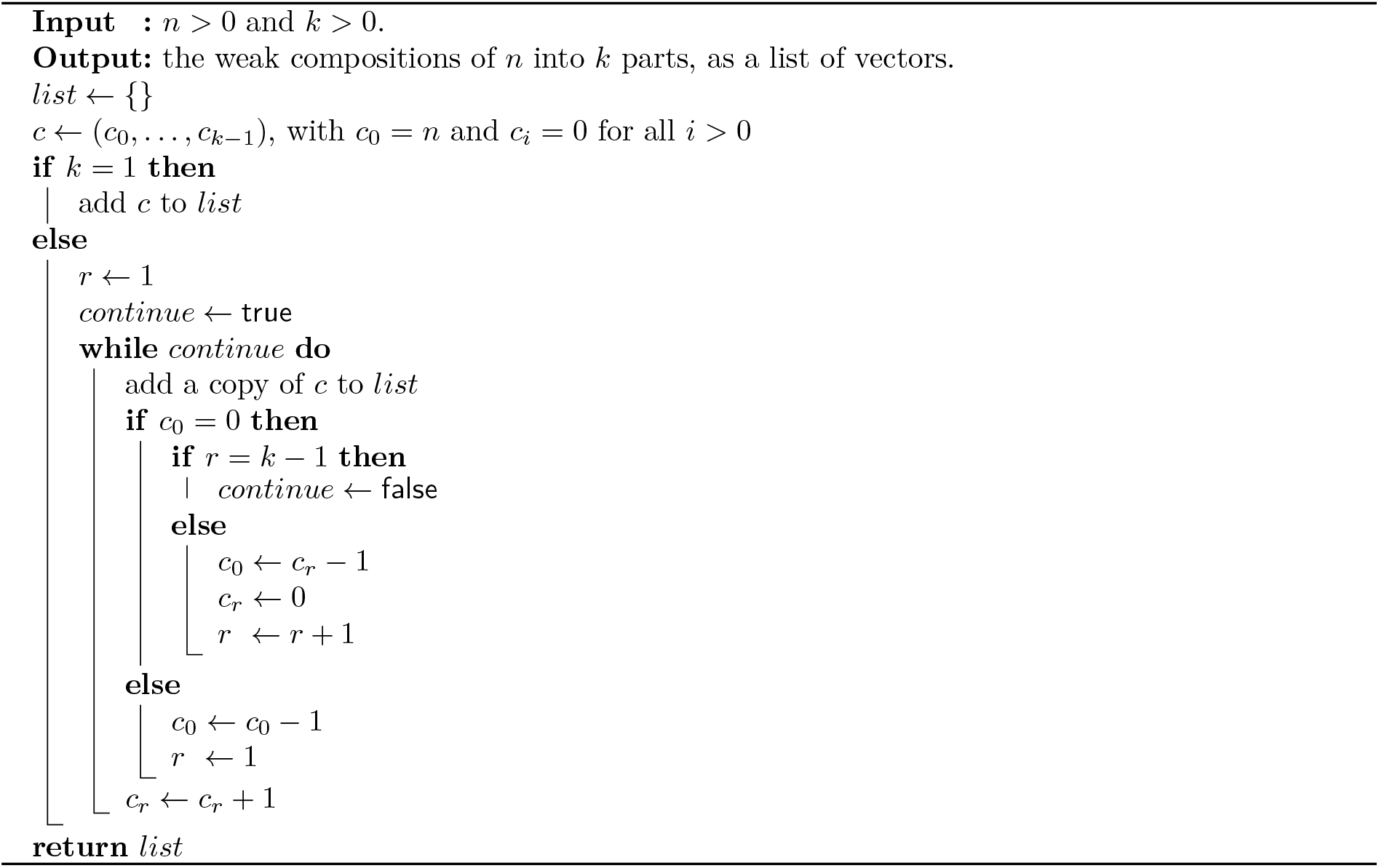

### SM.5 The previous-generation and Euler–Lotka matrices

The population projection matrix **A** projects the population one time step into the future, whereas the genealogical matrix **P** projects the lineage of an individual one time step into the past. Just as the so-called next-generation matrix **R** = **F**(**I** − **S**)^−1^ allows us to project the population one *generation* into the future (see e.g. Caswell (2001), Chapter 5), it is possible to define a matrix **G** that projects the lineage of an individual one generation into the past. We term this matrix the *previous-generation matrix*, and show that it has interesting connections with the Euler–Lotka equation.

Recalling the interpretation of the matrice **P**_**S**_ and **P**_**F**_ detailed in Section 2.1, we see that when going up the lineage of an individual *ego* from class *i* the probability that we reach its mother *t* time steps ago and in class *j* is the (*i, j*)-th entry of the matrix 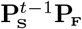. Considering all possible *t* at which this mother can be reached shows that the probability that the mother of *ego* was in class *j* when she gave birth to *ego* is the (*i, j*)-th entry of the matrix

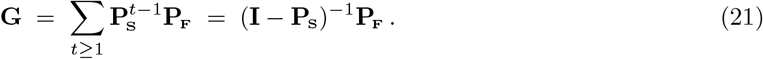

Note that (**I** − *P*_**S**_)^−1^ is always well-defined, and that it is the fundamental matrix of the absorbing Markov chain associated with **P**_**S**_ (see e.g. Kemeny and Snell (1976), Chapter V). The Markov chain associated with **G** is not ergodic (because non-reproductive classes are inaccessible), but its restriction to reproductive classes is. Thus, **G** has a stationary distribution ***ν*** that can be expressed as a function of the stationary distribution ***π*** of **P** (which, as we may recall, is itself given by the class reproductive values *π*_*i*_ = *v*_*i*_*w*_*i*_):

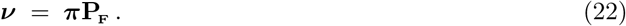

If this is not clear from the probabilistic interpretation, consider the following calculation:

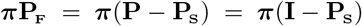

and therefore

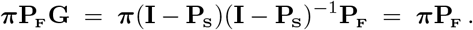

A notable fact about the previous-generation matrix **G** is that it corresponds to the transition matrix of the genealogical Markov chain associated with the projection matrix

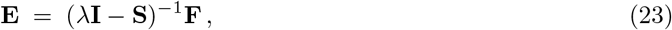

as proved in Proposition 3 below. Even though **E** is not primitive (because the non-reproductive classes of **A** correspond to “dead-ends” in the life-cycle graph associated to **E**), it has a dominant eigenvalue and a unique dominant right-eigenvector: 1 is an eigenvalue of **E** and it is greater in modulus than any of its other eigenvalues; and **w**, the stable class distribution of **A**, is the unique right-eigenvector of **E** associated with the eigenvalue 1 (see Proposition 1). Thus, since **E** can also be written

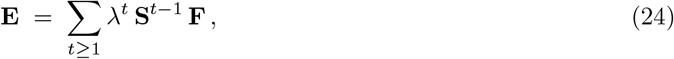

the equation **Ew** = **w** is a generalization of the Euler–Lotka equation to arbitrarily structured populations. Indeed, in age-based models, the Euler–Lotka equation (Lotka, 1939) is

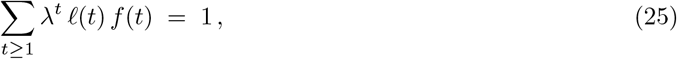

where *ℓ*(*t*) is the probability of surviving to age *t* and *f* (*t*) is the fertility at age *t*. But, in a general model: (1) the probability of surviving to age *t* depends both on the class in which the individual was born and on its class at age *t*, and it is given by the matrix **S**^*t*^; (2) the fertilities depend only on the class of the individual at age *t* (which may or may not include age as a variable) and are given by **F**.

It should be noted, however, that although **E** provides a very natural generalization of the Euler–Lotka equation, another natural generalization is possible. Indeed, consider the matrix

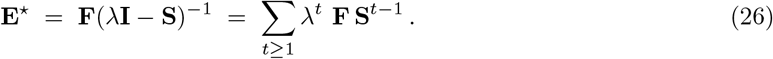

Then, as shown in Proposition 1, **E**\* also has dominant eigenvalue 1 and satisfies **vE**\* = **v**, where **v** is the vector of reproductive values of **A**. As a result, the equation **vE**\* = **v** can also be viewed as a natural generalization of the Euler–Lotka equation. Finally, note that **E**\* can be interpreted as the next-generation matrix associated with the projection matrix *λ*^−1^ **A**.

Because of their interpretation as what are arguably the two most natural generalizations of the Euler–Lotka equation, we term the matrices **E** and **E**\* the *Euler–Lotka matrices*.

We now close this section by proving our various claims about the matrices **G**, **E** and **E**\*.

#### Proposition 1.

*Let* **A** = **S** + **F** *be a primitive matrix with dominant eigenvalue λ and dominant left and right eigenvector* **v** *and* **w***, respectively. Assume that the matrix* **S** *is convergent and that* **F** ≠ **0***, and define the matrices*

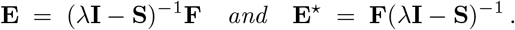

*Then,* **E** *and* **E**\* *have dominant eigenvalue 1 and*

i. **E w** = **w**.
ii. **v E***\** = **v**.

*Moreover, those are the only dominant eigenvectors (right or left, respectively) of those matrices.*

*Proof.* Let us start by proving (i) and (ii). First, note that

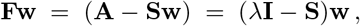

and therefore

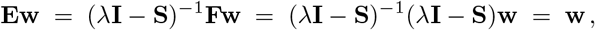

proving (i). Similarly, we get (ii) by noting that **vF** = **v**(*λ***I** − *S*).

Let us now prove that **E** and **E**\* have spectral radius 1. Let *ρ* denote the spectral radius. Note that since *ρ*(**AB**) = *ρ*(**BA**) for every matrices **A** and **B** (see e.g. Meyer, 2000, Exercice 7.1.19), we immediately have *ρ*(**E**) = *ρ*(**E**\*); moreover, from (i) and (ii), this spectral radius is at least 1.

Now, the “min-max” Collatz–Wielandt formula for non-negatives matrices (see Equation 8.3.3 in Meyer (2000) for the usual “max-min” version, and Exercice 8.2.9 for the “min-max” version) gives

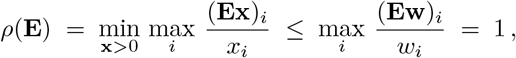

since **w** > 0 and **Ew** = **w**.

To conclude the proof, we have to show that **E**(resp. **E**\*) has no other eigenvalue with modulus 1, and that **w** (resp. **v**) is the only right-eigenvector (resp. left-eigenvector) associated with the eigenvalue 1. For this, consider the life-cycle graph associated with **E**, and note that non-reproductive classes (that is, *j* such that *f*_*ij*_ = 0 for all *i*) are terminal vertices in that graph. By contrast, the set of reproductive classes is irreducible (to find a directed paths between two vertices in the graph of **E**, consider the restriction of one such path in the graph of **A**– which is guaranteed to exists because **A** is irreducible – to the reproductive classes). The result then follows from SM.3. The matrix **E**\* is treated similarly, but using newborn classes (that is, *i* such that there exists *j* with *f*_*ij*_ > 0) instead of reproductive ones.

#### Remark 2.

The fact that *ρ*(**E***\**) = 1 can also be seen from the interpretation of **E**\* as the next-generation matrix of the projection matrix *λ−*1**A**: indeed, as such, *ρ*(**E**\*) is the *R*_0_ associated with the matrix *λ*^−1^**A**, whose asymptotic growth rate is 1.

#### Proposition 3.

*The previous-generation matrix* **G** = (**I**−**P**_**S**_)^−1^**P**_**F**_ *is the transition matrix of the genealogical Markov chain associated with the Euler–Lotka matrix* **E** = (*λ***I**−**S**)^−1^**F**.

*Proof.* First, note that since, by Proposition 1, **E** has dominant eigenvalue 1 and a unique dominant right-eigenvector **w**, its genealogical Markov chain is well-defined and given by Equation (1). Therefore, we only have to show that the entries of **G** and **E** satisfy:

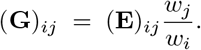

For this, let us note that, for all *t* ≥ 1,

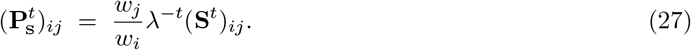

Indeed, the case *t* = 1 is the definition of **P**_**S**_ = (*p*_**S**_(*i, k*)), see Equation (2). Thus, by induction,

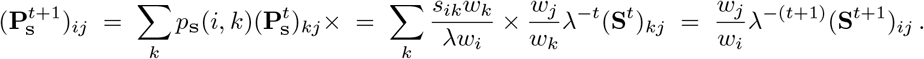

Now, from 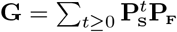 we get

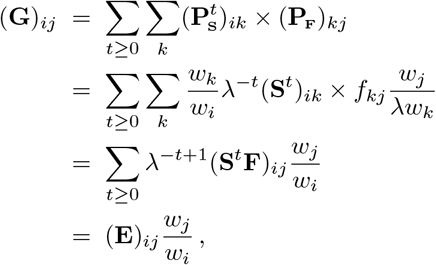

concluding the proof.

### SM.6 Upper bound on 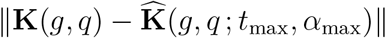

In this section, we provide an upper bound on the error that results from using a finite sum when computing the kinship matrix, as explained in Section 2.5. This upper bound can be used to determine values of *t*_max_ and *α*_max_ that ensure we meet a given accuracy in our estimation of **K**. It should be noted, however, that this upper bound is rather crude. As such, it will sometimes greatly overestimate the actual error – to the point of being uninformative. This is especially likely to happen in species with high survival and a low fertility.

Let || · || be any submultiplicative matrix norm, meaning that for any two matrices **A** and **B** we have ||**AB**|| ≤ ||**A**|| × ||**B**||. Note that most of the “standard” matrix norms (the induced norms, the Frobenius norm, the nuclear norm, etc) are submultiplicative, one notable exception being the max norm. Set

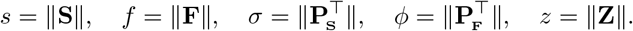

For our upper bound to be useful, the norm || · || should be chosen so that *sσ* < 1, ideally with *sσ* as small as possible. Note that the existence of such a norm is guaranteed by the fact that the matrices **S** and **P**_**S**_ are convergent (that is, **S***t* → **0** and 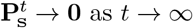). In particular, the *ℓ*_1_ operator norm, given by

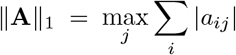

always satisfies 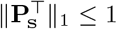 and ||**S**||_1_ < 1, unless there exists a class from which individuals cannot die. However, in practice we do not necessarily recommend sticking to this norm, but instead comparing several of them and choosing the one that minimizes *sσ*.

Once a norm has been chosen and the quantities *s*, *f*, *σ*, *ϕ* and *z* have been computed accordingly, our upper bound on 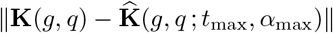 is

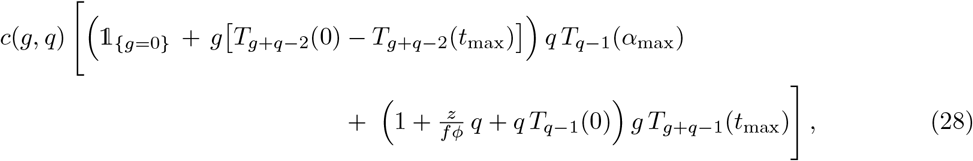

where

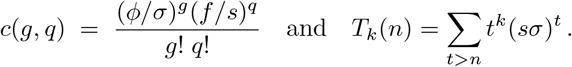

Note that this upper bound goes to zero as *t*_max_ and *α*_max_ go to infinity if and only if *sσ* < 1. To compute it numerically, *T* can be expressed using special functions; for instance,

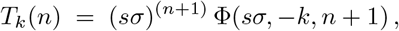

where Φ denotes the Lerch transcendent, which is available in many programming languages (Python, R, Mathematica, etc). In languages that do not implement the Lerch transcendent, such as MATLAB, the polylogarithm can be used instead:

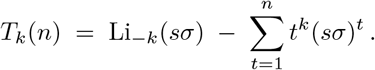

It is also relatively straightforward to design an ad hoc implementation of *T*.

The rest of this section is devoted to the calculation of the upper bound given in (28). Let us start by introducing some notation. As in the rest of this manuscript, we use the convention 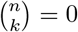 when *n <* 0 or *k* ∉ {0, …, *n*}, so that we always have

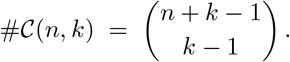

We also use the habitual convention 1*/k*! = 0 when *k* = −1, −2, …

#### Lemma 4.

i. *For g* = 0, ||**U**(0, 0)^⊤^|| = 1 *and* ||**U**(*t,* 0)^⊤^|| = 0 *when t >* 0.
ii. *For g* ≥ 1*, and t* ≥ 0, 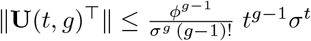.
iii. *For t* ≥ 2 *and any q,* 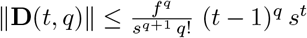.
iv. *For t* ≥ 0 *and any q,* 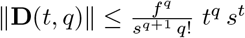.

*Proof.* Point (i) is immediate, since **U**(*t,* 0) = **I** if *t* = 0 and **0** otherwise, and ||**I**|| = 1 for any submultiplicative matrix norm.

For point (ii), using 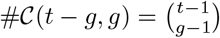 and the submultiplicativity of the norm gives

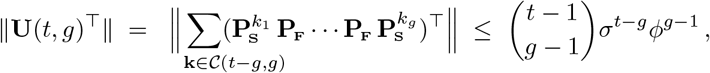

and the result follows from the inequality 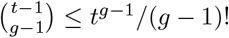. Similarly,

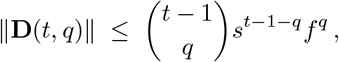

proving (iii) and (iv) since 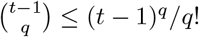 for *t* ≥ 2 and 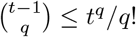 for *t* ≥ 0.

*Proof of Inequality 28.* With the notation of Section 2, let us write

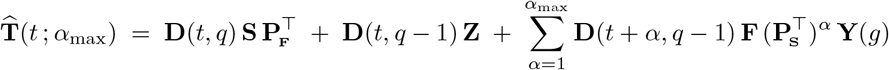

and 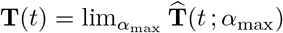. Note that we have kept the dependency on *g* and *q* implicit to ease the notation, and will allow ourselves to do so with other matrices such as **K** and **U** when there is no risk of confusion. With this notation, we have

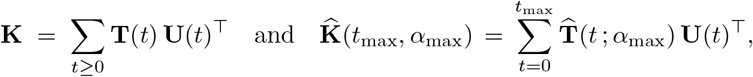

and, therefore,

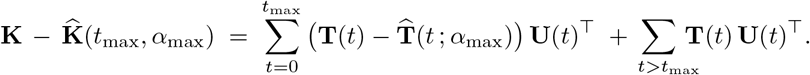

Using, again, the submultiplicativity of the norm, this yields

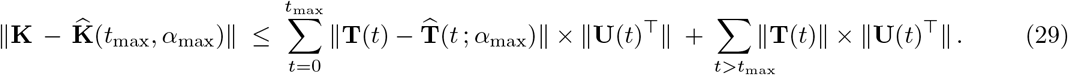

Let us assume that *g* ≥ 1 and treat the case *g* = 0 zero separately later. Then, 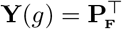 and therefore

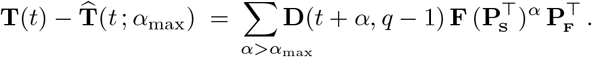

Note that since **U**(0) = **0**, we can have *t* ≥ 1 and *α* ≥ 1 in the sum. As a result, we can apply point (iii) of Lemma 4 to **D**(*t* + *α, q* − 1) and get

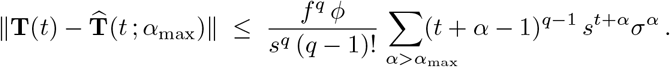

Using that (*t* + *α* − 1)^*q*−1^ ≤ *t*^*q−*1^*α*^*q*−1^ for all *t* ≥ 1 and *α* ≥ 1 then gives

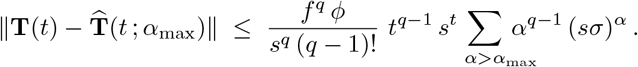

Using point (ii) of Lemma 4 in this expression, we get

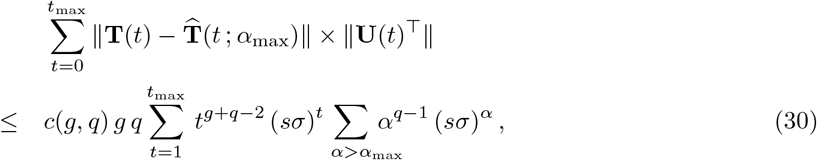

where

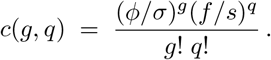

Similarly, applying Lemma 4 in

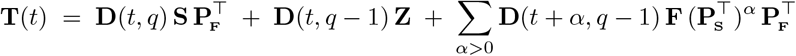

yields, after some rearranging and using that *t*^*q*−1^ ≤ *t*^*q*^,

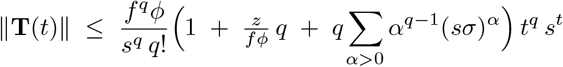

Therefore,

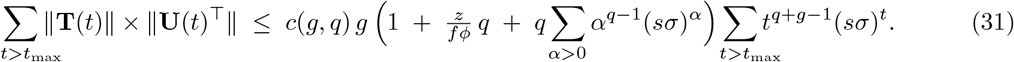

Plugging Inequalities (30) and (31) in (29) gives us the following upper bound for *g* ≥ 1

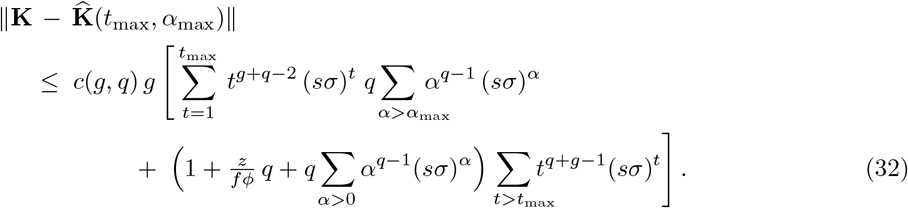

Note that, because of the factor *g*, this upper bound is equal to 0 when *g* = 0.

Let us now turn to the case *g* = 0. In that case, the expression of the kinship matrix reduces to

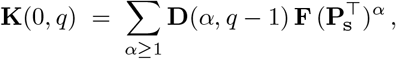

see Equation (12) in the main text. Therefore,

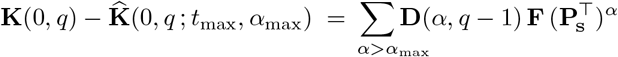

and using point (iv) of Lemma 4 we get

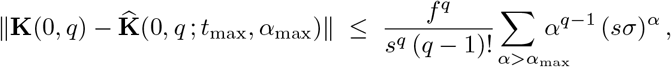

which, since *c*(0*, q*) = (*f/s*)^*q*^/(*q* − 1)!, can also be written

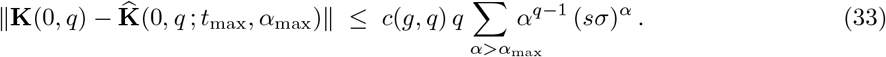

Combining (32) and (33), we get the following general upper bound

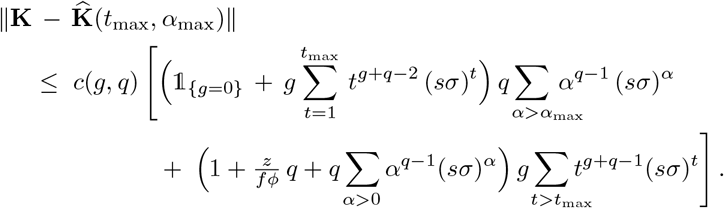

Letting

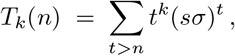

the right-hand side of this inequality can also be written

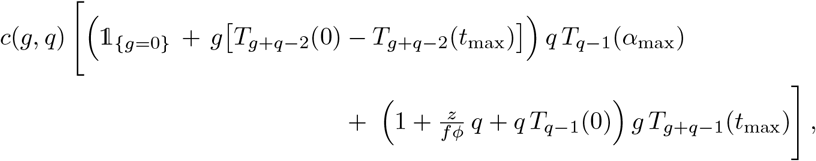

yielding the upper bound (28).

### SM.7 Capping age in a model

In this section, we briefly describe a straightforward way to turn a model without a maximal age (that is, such that **S**^*t*^ ≠ 0 for all *t*) into a model with maximal age *ω* (that is, where **S**^*ω*−1^ ≠ **0** and **S**^*ω*^ = **0**). This could be useful, e.g, to estimate the precision of the calculation of the kinship matrix done by choosing *t*_max_ and *α*_max_ as a function of *ω*, as discussed in Section 2.5. However, there are some caveats associated with this “capping of age”, as the resulting model is non-trivially related to the original one.

First, given a projection matrix **A** = **S** + **F**, choose a maximal age 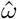 (meaning that individuals will *not* reach age 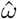 in the capped model). This could be done:

- either using some a priori knowledge on the biology of the species / population under study;
- or from the model itself, by picking *ε* such that the probability of reaching age 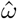 is less than *ε*. This can be done as follows: letting **x** = (*x*_*i*_) be defined by *x*_*i*_ = 1 if *i* is a newborn class and *x*_*i*_ = 0 otherwise, we have: 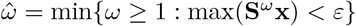.

Once 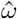 has been chosen, one way to obtain a projection matrix with maximal age 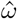 is the following. Start by considering the matrix 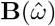 defined, using block-notation, by

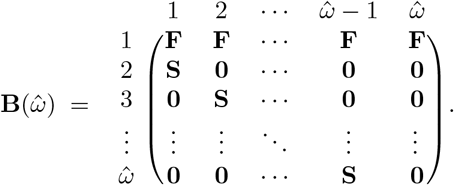

Such “block-Leslie” matrices are frequently used to convert *stage* classified models into *age × stage* classified models – see for instance the matrix **M**_AS_(*p*) of Lebreton (2005), which is almost identical to this one with the exception of the 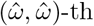 block, which we take to be **0** whereas Lebreton takes it to be **S**.

Note that the matrix 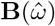 is aperiodic, but that it is *not* irreducible. For this reason, it will be more convenient to work with the matrix 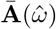 obtained by keeping only the classes of 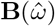 that have a non-zero asymptotic abundance, as explained in SM.3.

Because the projection matrix 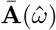 is primitive and has a maximal age 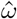, its kinship matrix can be computed exactly by taking 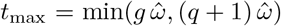 and 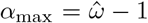 in Equation (15). This yields a kinship matrix with the same dimensions as 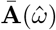. This kinship matrix 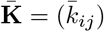 can be turned back into an *n × n* kinship matrix, where *n* is the dimension of **A**, by summing the entries that correspond to the same class of **A** while weighting each entry 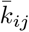 by the asymptotic abundance 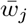.

However, some care is needed when interpreting the resulting kinship matrix: indeed, the capped matrix 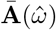 describes different population dynamics than the matrix **A**; for instance, it has a smaller asymptotic growth rate, a shorter generation time, etc. As a result, it may or may not better describe the population under study than the matrix **A**, and it is the responsibility of the ecologist to determine whether this is the case.

Finally, note that if all descriptors of **A** and 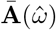 are very similar, this tends to suggest that neglecting individuals that reach age 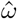 has little influence on the descriptors of the model, and could thus indicate that the kinship matrices obtained from **A** by taking 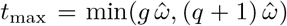 and 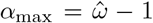 in Equation (15) are fairly accurate. This gives a way to get a rough idea of the precision of the estimate 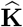 when the upper bound of SM.6 is not useful.

### SM.8 Expression of K(*g, q*) for 1×1 models

Assume that the projection matrix is a 1×1 matrix **A** = (*s* + *f*), with *f* > 0 and 0 ≤ *s* < 1, and let *λ* = *f* + *s*. In that case, the genealogical matrices **P**_**S**_and **P**_**F**_, given by Equation (2) in the main text, are

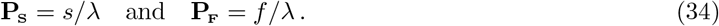

Since these matrices commute and that 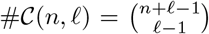, the expression of the matrix **U**(*t, g*) given by Equation (4) becomes: **U**(0, 0) = 1 and, for any other values of *t* and *g*,

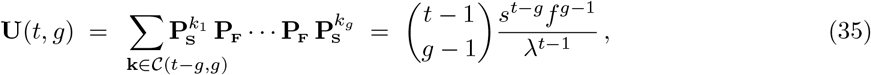

with the convention that 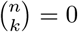 when *n* < 0 or *k* ∉ {0, …, *n*}. Similarly, Equation (5) gives

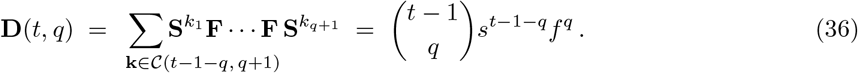

Finally, the same-litter newborn sisters matrix **Z** (see 2.4) can be written 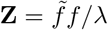, where 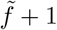 is the expected number of offspring conditional on reproduction.

Plugging all of these quantities in Equation (13), we get the expression of **K** for 1×1 models. First, for *g* = 0,

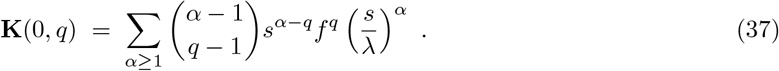

Second, for *g* ≥ 1,

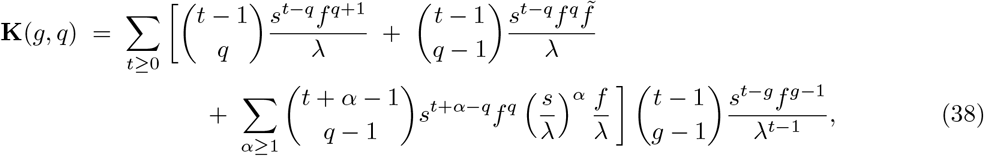

again with the convention that 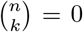 when *n <* 0 or *k* ∉ {0, …, *n*}. Rearranging a bit, this second expression can also be written

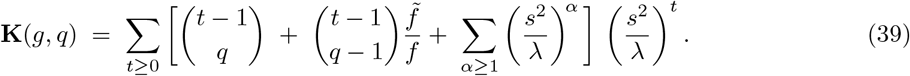

### SM.9 Asymptotic behavior of 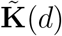 as d → ∞

In this section, we detail the mathematical reasoning between the introduction and characterization of the statistic *θ*_c_ discussed in Section 2.6. Loosely speaking, *θ*_c_ is such that, for any matrix norm,

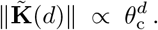

More rigorously, 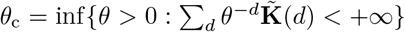. This justifies the appellation *exponential growth rate of* 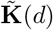. Our main result is that *θ*_c_ admits the following characterization, given as Equation (18) in the main text:

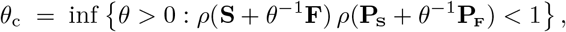

where *ρ* denotes the spectral radius (here, the dominant eigenvalue, since the matrices are primitive). This characterization makes it straightforward to compute *θ*_c_ numerically (e.g, using the bisection method, as implemented in our code, see SM.1).

Although the result for 1×1 models follows as a special case of the result for *n*×*n* models, we start by treating it on its own, in full detail. This is because the expression of **K**(*g, q*) for 1×1 models given in SM.8 and the fact that we are dealing with numerical (instead of matricial) series make the discussion less technical, while retaining its main idea; it is thus a useful stepping-stone for the *n*×*n* case. In addition to this, the simple expression of *θ*_c_ for 1×1 models deserves a special discussion.

#### SM.9.1 Expression of *θ*_c_ in 1×1 models

As in SM.8, assume that the projection matrix is a 1×1 matrix **A** = (*s* + *f*), with *f* > 0 and 0 ≤ *s* < 1, and write *λ* = *f* + *s*. For any *θ >* 0, let us consider the sum

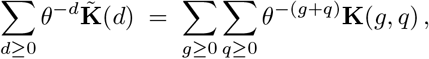

this equality being true because the summands are non-negative and thus the order of summation is irrelevant (Fubini-Tonelli theorem). Our aim is to get a necessary and sufficient condition on *θ* for this series to be convergent.

Plugging Equations (37) and (38) into 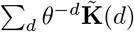, then using again Fubini-Tonelli’s theorem to
change the order of summation and rearranging a bit, we get

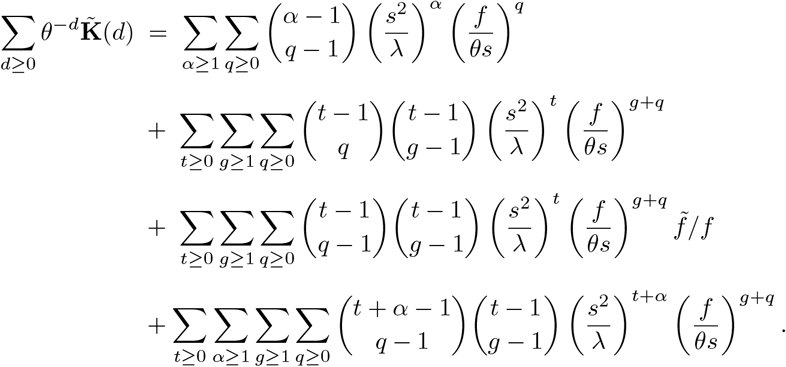

Let us start by focusing on the first term of the right-hand side. Recalling that we use the convention 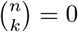 when *n* < 0 or *k* ∉ {0, …, *n*} and using the binomial theorem, we have, for every *α* ≥ 1,

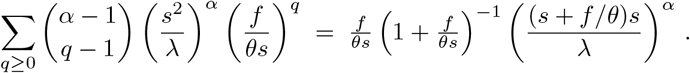

Therefore,

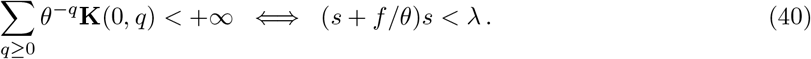

Finally, since *λ* = *s* + *f*,

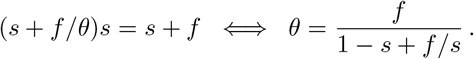

This means that, as *q* increases, the number of *q*-th generation descendants of *ego* grows roughly like *γ*^*q*^, where 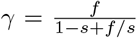. Unsurprisingly, this exponential rate of increase *γ* is exactly the expected number of daughters of *ego* that are currently alive, that is,

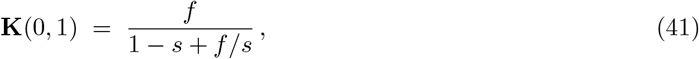

as can be checked from Equation (37).

Let us now turn to the second term of the right-hand side of the expression of 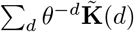. Again, for every *t* ≥ 1,

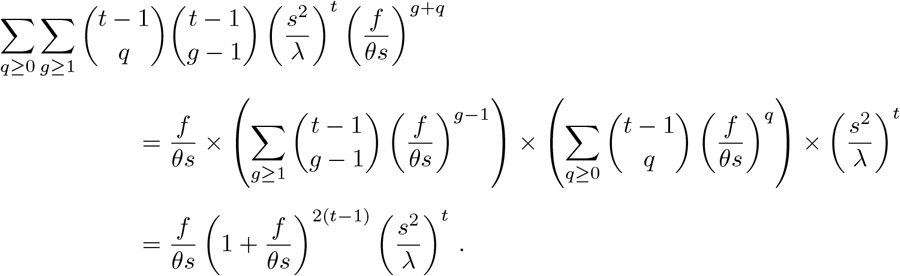

Therefore,

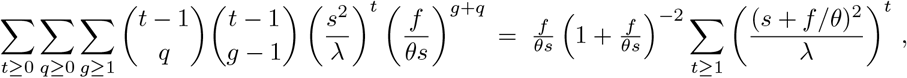

which converges if and only if (*s* + *f/θ*)^2^ < *λ*. Similarly,

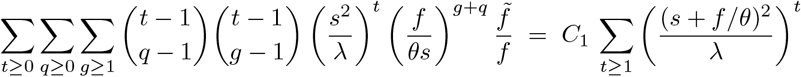

for some irrelevant constant *C*_1_, and

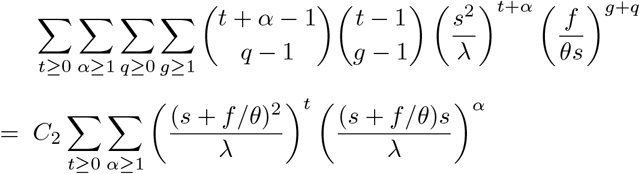

for some constant *C*_2_. This last series converges if and only if (*s* + *f/θ*)^2^ < *λ* and (*s* + *f*/*θ*)*s* < *λ*; but, since *s < s* + *fθ*, these two conditions are equivalent to (*s* + *f*/*θ*)^2^ < *λ*.

Putting the pieces together, we see that

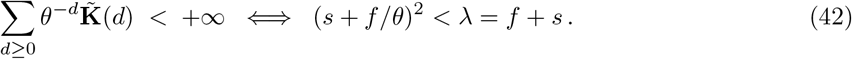

Again, loosely speaking, this means that 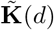 grows like 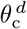, where

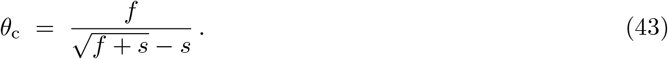

(note that *θ*_c_is always well-defined and positive because *f* > 0 and 0 ≤ *s* < 1). Interestingly, whenever *s* ≪ *f*, be it because *s* ≪ 1 or because *f* ≫ 1, 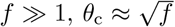 does not depend on *s*. Also note that *θ*_c_ can be written

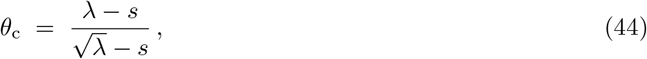

which is useful because it shows that:

i. As expected, *θ*_c_ > 1 if and only if *λ >* 1.
ii. For a fixed value of *λ*, *θ*_c_ is an increasing function of *s* when *λ* > 1 and a decreasing function of *s* when *λ* < 1.

In particular, the second point reflects the intuitive fact that, if there is an advantage to having closely related kin, then given two mutants with the same growth rate, the mutant with the higher survival is expected to be favored – in other words, that *K* strategies are expected to evolve when close kin help each other. What is perhaps surprising is that this tendency is reversed when the population is decreasing. It should be noted, however, that our setting assumes that the population has been at the demographic steady state for a long period of time, which undermines the relevance of our formulas when *λ* < 1.

#### SM.9.2 Characterization of *θ*_c_ in *n*×*n* models

Despite additional technicalities, the case of *n*×*n* models is very similar to that of 1×1 ones. Here the crucial observation is that

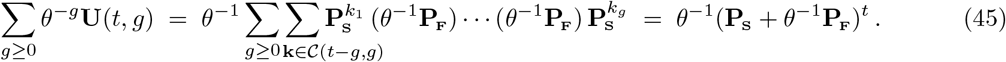

Similarly,

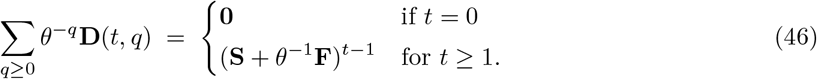

Let us focus on the growth of the term **K**_y_ in **K** = **K**_y_ + **K**_s_ + **K**_o_, that is,

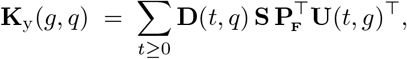

as given by Equation (6). Using the same reasoning as for 1×1 models, but with Equations (45) and (46) instead of the binomial theorem, we get

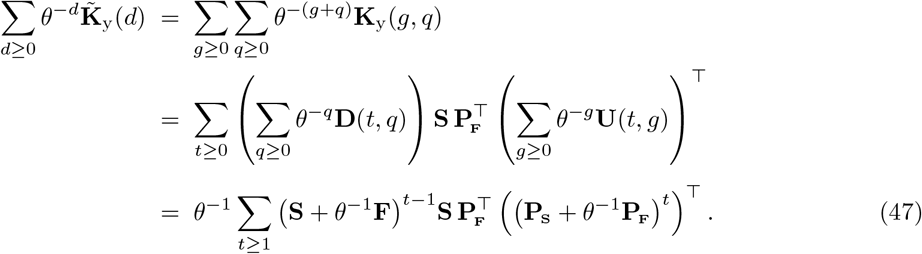

Now, observe that the matrices **A**_*θ*_ = **S** + *θ*^−1^**F** and **P**_*θ*_ = **P**_**S**_ + *θ*^−1^**P**_**F**_ are both primitive (because **A** = **S** + **F**, and thereby **P** = **P**_**S**_ + **P**_**F**_, are). As a result, letting *ρ*(·) denote their dominant eigenvalue, as *t* → ∞,

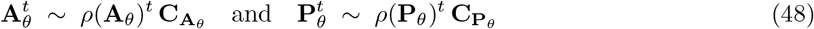

for some irrelevant matrices 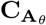 and 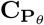. Combining this with (47), we get

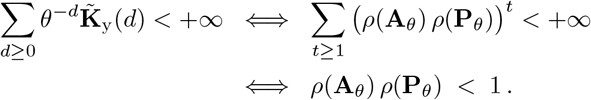

The growth of the matrices **K**_s_ and **K**_o_ is treated similarly and yields the same criterion for convergence, completing our charaterization of *θ*_*c*_.

Note that if **A** = (*s* + *f*) is a 1×1 matrix, then *ρ*(**A**_*θ*_) = *s* + *f/θ* and 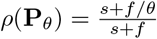 and so we fallback on Equation (42).

Finally, because *f* : *θ → ρ*(**A**_*θ*_)*ρ*(**P**_*θ*_) is increasing in *θ* and easily evaluable numerically, this makes the numerical determination of *θ*_*c*_ straightforward and efficient. Also note that to get an initial estimate of *θ*_*c*_ it si possible to “collapse” **A** into a 1×1 model, as explained in Bienvenu et al. (2017), and then to use Equation (44). Using individualistic collapsing yields

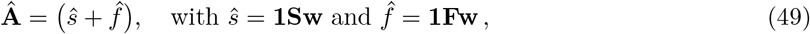

where **1** = (1, …, 1), whereas using genealogical collapsing – which is expected to be more accurate for the estimation of most biological descriptors – yields

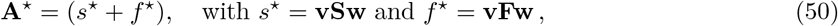

where the vector of reproductive values **v** is scaled in such a way that **vw** = 1.

### SM.10 Computing time of 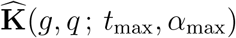

The computing time of the matrix 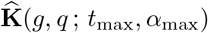, as given by Equation (15), increases rapidly as a function of *g*, *q*, *t*_max_ and *α*_max_, and can be prohibitively long in practice. To help assess this, in this section we provide rough estimates of the time complexity of a standard implementation of 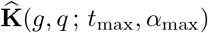.

#### SM.10.1 Setting and notation

We denote by 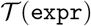 the number of elementary operations required to compute the expression expr, where by “elementary operation” we refer to: matrix addition; matrix multiplication; matrix transposition; and matrix power. All of these operations are assumed to a have a Θ(1) time complexity, independently of the size *n* of the matrix, which we assume to be fixed. Should large matrices be studied, this Θ(1) could be replaced by the complexity of the matrix product, that is, Θ(*n*^*x*^) for some *x ∈*[2, 3].

In reality, the time complexity of the matrix power **A**^*k*^ is Θ(log *k*); but since taking this into account has the effect of adding a logarithmic factor Θ(log(*t*_max_ + *α*_max_)) and that, for all practical purposes, this factor can be assumed to be bounded, we use Θ(1) for simplicity.

Finally, to alleviate the notation, we write *f*(*t*_max_*, α*_max_) ≍ expr to indicate that *f* (*t*_max_*, α*_max_) = Θ(expr) as *t*_max_ and *α*_max_ increase. Note that because we are interested in what happens not only when *g* and *q* are fixed but also when they increase with *t*_max_ and *α*_max_, we will implicitly treat them as functions of *t*_max_ and *α*_max_, rather than as constants.

#### SM.10.2 Calculation

First, observe that, since 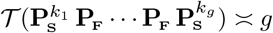 and 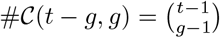, for *g* ≥ 1 we have

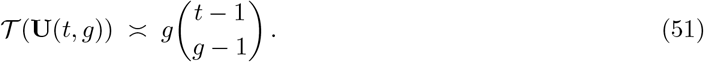

Similarly, for *q* ≥ 1,

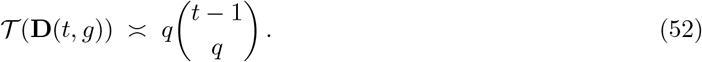

Now, recall Equation (15):

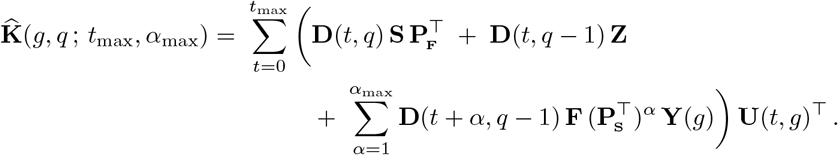

Focusing on the case *g* ≥ 1*, q* ≥ 1, we get

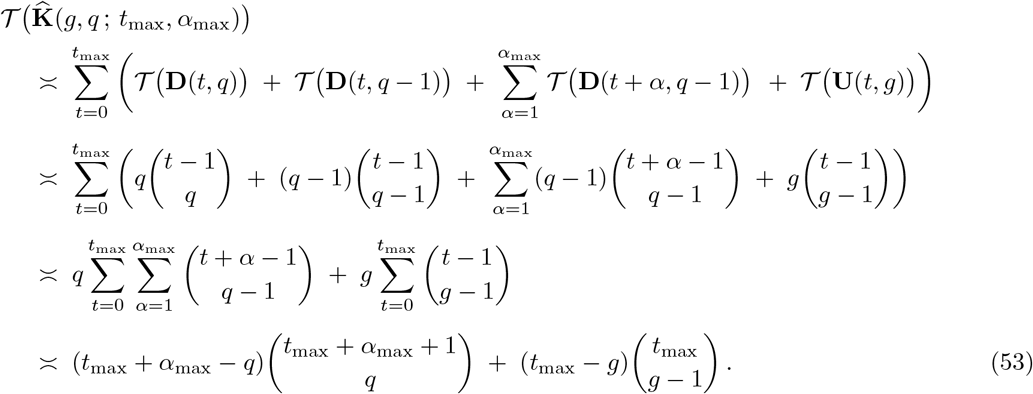

Let us focus on two specific cases: first, assume that *g* and *q* are constant. This is the most relevant scenario for practical applications, where *g* and *q* will typically be very small. In that case, as *t*_max_ and *α*_max_ go to infinity,

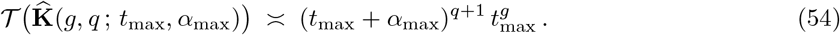

In particular, if *t*_max_ and *α*_max_ are chosen according to Equation (16), as a function of a maximal age *ω* (be it a “true” maximal age built in the model, or an inferred one used in the computation of 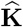, then

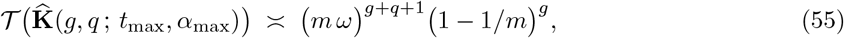

with *m* = min(*g, q*+1)+1. Assuming that the computing time start being prohibitive after 10^9^ elementary operations (which, of course, is going to be highly dependent on the specificities of the implementation of 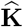 and on the computing power available), Table 2 below gives the maximal value of *ω* for which the calculations can be performed as a function of *g* and *q*, based on Equation (55).

**Table 2:** Largest value of *ω* for which (*m ω*)^*g*+*q*+1^(1−1*/m*)^*g*^ < 10^9^, with *m* = min(*g, q* + 1) + 1. This gives a rough idea of the values of *g* and *q* for which **K**(*g, q*) can be estimated, as function of the maximal age *ω* in the life cycle. More precise estimations can be obtained, e.g., using Equation (53) or even one of the more precise estimates that precede it.

| $q \backslash g$ | 0 | 1 | 2 | 3 | 4 | 5 | 6 | 7 |
| --- | --- | --- | --- | --- | --- | --- | --- | --- |
| 0 | — | 31622 | 999 | 177 | 63 | 31 | 19 | 13 |
| 1 | 22360 | 629 | 105 | 36 | 17 | 10 | 7 | 5 |
| 2 | 793 | 72 | 24 | 12 | 7 | 4 | 3 | 2 |
| 3 | 149 | 26 | 9 | 5 | 3 | 2 | 2 | 1 |
| 4 | 54 | 13 | 5 | 2 | 2 | 1 | 1 | 1 |
| 5 | 28 | 8 | 3 | 2 | 1 | 1 | 1 | 0 |
| 6 | 17 | 6 | 3 | 1 | 1 | 0 | 0 | 0 |
| 7 | 12 | 4 | 2 | 1 | 1 | 0 | 0 | 0 |

Finally, let us focus on the worst-case scenario and assume that *q* = ⌊(*t*_max_ + *α*_max_)*/*2⌋. In that case, Equation (53) gives

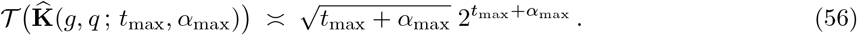

Of course, the same reasoning shows that the time complexity is 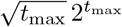 when *g* = ⌊*t*_max_/2⌋. This shows that there is no hope to be able to compute **K**(*g, q*) for large values of *g* and/or *q*. However, note that some information about the large-scale structure of the kinship can still be obtained through the statistics *θ*_c_ discussed in SM.9.

### SM.11 Matrilineal kinship and relatedness

In this section, we briefly explain why – under some fairly stringent hypotheses – the matrilineal notion of kinship that we consider (that is, the expected of kin of *ego* in the genealogical tree of females) coincides with the usual notion of relatedness in population genetics (that is, the expected number of genes of *ego* that are identical by descent with those of its kin, including males) – and why it can, under more acceptable hypotheses, be seen as a lower bound on relatedness.

Consider the idealized setting of an infinite panmictic population where there is no difference in survival or reproduction between males and females, and where exactly one offspring is produced at each reproductive event (so that no two individuals share the same two parents). In that setting, *ego* has exactly 2^*g*^ distinct *g*-th generation ancestors in its complete pedigree – as opposed to one through matrilines only. Because the population is infinite and panmictic, these ancestors do not have shared descendants. Thus, the number of individuals that are *q*-th generation descendants of at least one of the *g*-th ancestors of *ego* is on average 2^*g*^ times the expected number of *q*-th descendants of any of those ancestors. Now, since there is no difference in survival and reproduction between males and females, the expected number of *q*-th descendants of an individual is 2^*q*^ times its number of *q*-th descendants through matrilines only. As a result, the expected number of (*g, q*)-kin in the complete pedigree of *ego* is 2^*g*+*q*^ times its number of matrilineal (*g, q*)-kin. Finally, since the relatedness of each of these (*g, q*)-kin with *ego* is 2^−(*g*+*q*)^, we see that the total amount of relatedness to *ego* that comes from its (*g, q*)-kin is, on average, exactly its number of matrilineal kin.

Of course, none of the assumptions on which this reasoning relies hold in the real world: populations are neither infinite nor panmictic, and a pair of individuals can give birth to more than one offspring – so that the pedigrees of the (*g, q*)-kin of *ego* will overlap. However, this overlap in itself is not a problem, as long as the expected number of descendants of each individual is the same. Indeed, assume for instance that two of the 2^*g*^ *g*-th ancestors of *ego* are, in fact, the same individual. Then, the contribution of those ancestors to the (*g, q*)-kin of *ego* is divided by two, but this is compensated by the fact that each of these individuals is twice as related to *ego* as if the two *g*-th ancestors had been different individuals. This reasoning can be generalized to the whole portion of the pedigree required to find the (*g, q*)-kin of *ego*.

Another consequence of the fact that the population is not infinite and panmictic is that the *g*-th ancestors of *ego*, even if they are all distinct, are bound to have some degree of relatedness that should be taken into account in an exact calculation of the relatedness, whereas we are ignoring it here. In other words, neglecting the relatedness of the least common ancestors of *ego* and its (*g, q*)-kin means that we tend to underestimate the true relatedness, and this effect is more pronounced in consanguineous populations.

Finally, the effect of a difference in survival between males and females is more complex, and is bound to non-trivially affect the number of kin of *ego*. However, if the sex-ratio is 1:1, then for this to remain constant over time males and females should have, on average, the same number of offspring across the course of their life. Therefore, even if the distribution of the number of (*g, q*)-kin of *ego* is modified, its expected value should not change – and thus neither does their contribution to the relatedness.

Altogether, these considerations suggest that the number of matrilineal kin computed by our formulas can be seen as a direct measure of relatedness in population with low consanguinity, and as a lower bound on relatedness in more consanguineous populations.

### SM.12 Comparison to individual-based simulations for the ground squirrel *Spermophilus dauricus*

To illustrate the validity of our formulas, we compare them to kinship matrices estimated from individual-based simulations. For this, we use the life-cycle of the ground squirrel *Spermophilus dauricus* presented in Box 2.

The simulations were done in R using the package NetLogoR (Bauduin et al., 2019), and correspond to a female-based population. At each time step, the individuals went through the following events, successively:

- **Reproduction:** the number of newborns produced by each female in stage *i* is drawn from a Poisson distribution with parameter *m*_*i*_, where *m*_*i*_ is the expected number of (female) newborns produced by each (female) individual. The newborns are temporarily put in an auxiliary stage *i* = 0 that does not correspond to a class of the matrix population model.
- **Survival:** each individual in stage *i* survives with probability *s*_*i*_, independently of the other individuals, and enters stage *i* + 1 (individuals in stage *i* = 3 remained in that stage). That is, conditional on the number *n*_*i*_ of individuals in stage *i* before the event, the number of individuals in stage *i* + 1 after the event is a binomial variable with parameters (*n*_*i*_, *s*_*i*_).

To run the simulations, we chose a maximum number of time-steps ensuring that, with high probability, for the values of (*g, q*) considered the simulated genealogy would contain the *g*-th and *q*-th ancestor of every individual alive at the end of the simulation. Here, using *T*_max_ = 50 times steps was sufficient.

Next, we chose an initial population ensuring that, with high probability, the population would not go extinct before *T*_max_. Here we started with 1010 individuals at the stable stage distribution, that is, with an initial population vector **n** = (470, 270, 270).

Finally, we ran 300 independent replicates of the population process. For each of those replicates, at the last time step, we computed the empirical kinship matrices for mothers and aunts:

- **K**_emp_(1, 0), whose (*i, j*)-th entry is fraction of individuals from class *j* whose mother is currently alive in stage *i*.
- **K**_emp_(2, 1), whose (*i, j*)-th entry is per-capita number of aunts alive in stage *i* for individuals in class *j*.

We averaged the empirical kinship matrices over all 300 replicates and compared their average 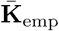 to the expected value **K** given by the formulas of the main text. For mothers, we have

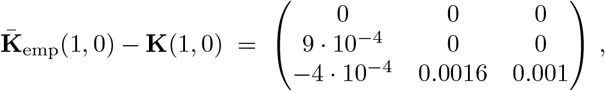

and for aunts we have

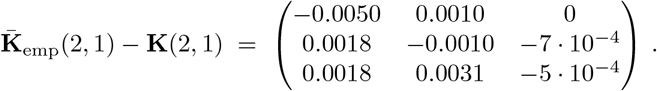

The empirical distribution of the sum of the columns of these matrices (that is the probability that the mother of *ego* is alive and its expected number of aunts, structured by the stage of *ego*) are depicted in Figures 5 and 6.

**Figure 5:**
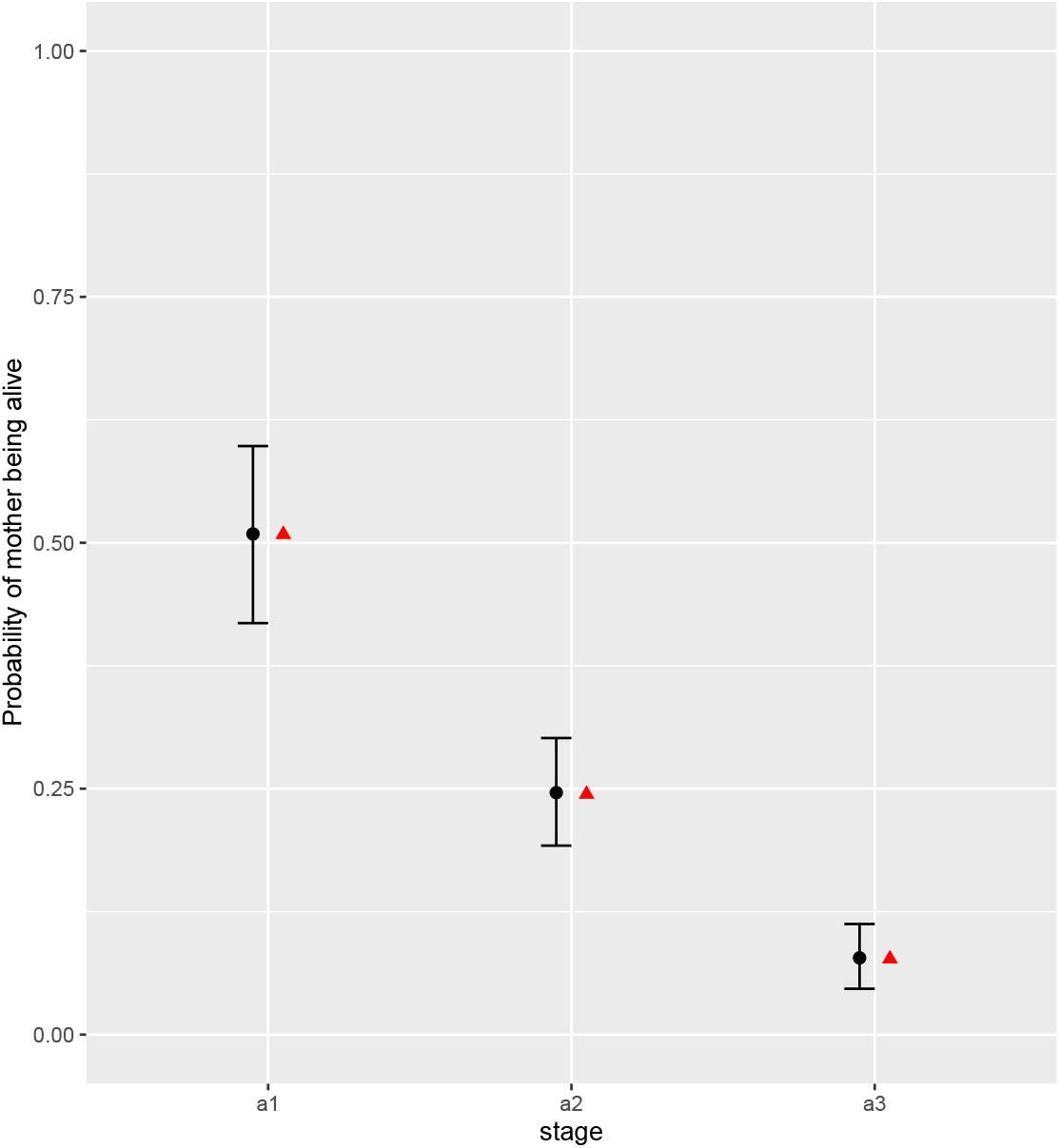
The empirical distribution of the probability that the mother of *ego* is alive, structured by the stage of *ego*. The black segments correspond to the 95% quantiles of the empirical distribution, the back dots to the empirical average, and the red triangles to the theoretical expected values given by our formulas.

**Figure 6:**
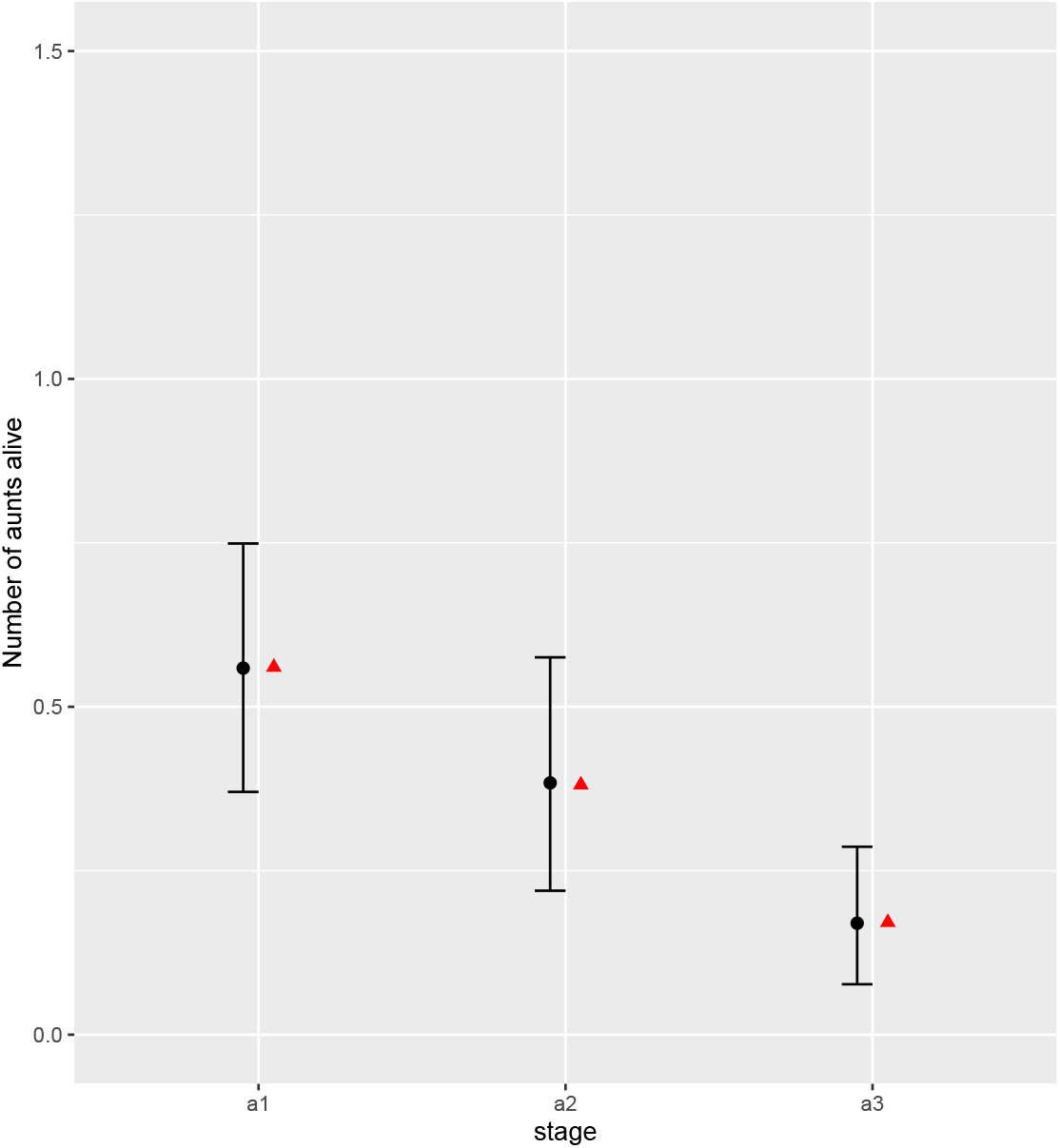
The empirical distribution of the average number of aunts of *ego*, structured by the stage of *ego*. The black segments correspond to the 95% quantiles of the empirical distribution, the back dots to the empirical average, and the red triangles to the theoretical expected values given by our formulas.

The code used in this Supplementary Material and the full output of the simulations can be found at https://github.com/SCubaynes/Squirrel_IBM.git.

## References

Atkins, J. R. (1974). On the fundamental consanguineal numbers and their structural basis. American Ethnologist, 1(1):1–31.

Barrai, I., Cavalli-Sforza, L., and Moroni, A. (1962). Frequencies of pedigrees of consanguineous marriages and mating structure of the population. Annals of Human Genetics, 25(4):347–377.

Bauduin, S., McIntire, E. J. B., and Chubaty, A. M. (2019). NetLogoR: a package to build and run spatially explicit agent-based models in R. Ecography, 42(11):1841–1849.

Bienvenu, F., Akçay, E., Legendre, S., and McCandlish, D. M. (2017). The genealogical decomposition of a matrix population model with applications to the aggregation of stages. Theoretical Population Biology, 115:69–80.

Bienvenu, F. and Legendre, S. (2015). A new approach to the generation time in matrix population models. The American Naturalist, 185(6):834–843.

Brault, S. and Caswell, H. (1993). Pod-specific demography of killer whales (Orcinus orca). Ecology, 74(5):1444–1454.

Caswell, H. (2001). Matrix Population Models. Sinauer Associates Inc., Sunderland, MA.

Caswell, H. (2019). The formal demography of kinship: A matrix formulation. Demographic Research, 41:679–712.

Caswell, H. (2020). The formal demography of kinship II: Multistate models, parity, and sibship. Demographic Research, 42:1097–1146.

Caswell, H. and Song, X. (2021). The formal demography of kinship iii: Kinship dynamics with time-varying demographic rates. bioRxiv.

Coresh, J. and Goldman, N. (1988). The effect of variability in the fertility schedule on numbers of kin. Mathematical Population Studies, 1(2):137–156.

Coste, C. F. D., Austerlitz, F., and Pavard, S. (2017). Trait level analysis of multitrait population projection matrices. Theoretical Population Biology, 116:47–58.

Coste, C. F. D. and Pavard, S. (2020). Analysis of a multitrait population projection matrix reveals the evolutionary and demographic effects of a life history trade-off. Ecological Modelling, 418:108915.

Cushing, J. M. and Zhou, Y. (1994). The net reproductive value and stability in matrix population models. Natural Resource Modeling, 8(4):297–333.

Demetrius, L. (1974). Demographic parameters and natural selection. Proceedings of the National Academy of Sciences, 71(12):4645–4647.

Demetrius, L. (1975). Natural selection and age structured populations. Genetics, 79(3):535–544.

Everett, C. J. and Ulam, S. (1948). Multiplicative systems, I. Proceedings of the National Academy of Sciences, 34(8):403–405.

Goodman, L. A., Keyfitz, N., and Pullum, T. W. (1974). Family formation and the frequency of various kinship relationships. Theoretical Population Biology, 5(1):1–27.

Hajnal, J. (1963). Concepts of random mating and the frequency of consanguineous marriages. Proceedings of the Royal Society of London. Series B. Biological Sciences, 159(974):125–177.

Joffe, A. and Waugh, W. A. O. (1985). The kin number problem in a multitype Galton-Watson population. Journal of Applied Probability, 22(1):37–47.

Kemeny, J. G. and Snell, J. L. (1976). Finite Markov chains. Springer-Verlag, New York, 2nd edition.

Knuth, D. E. (2011). Art of Computer Programming, Volume 4A. Addison-Wesley Professional.

Le Bras, H. (1973). Parents, grands-parents bisaieux. Population (French Edition), 1(1):9–38.

Lebreton, J.-D. (1990). Modelling density dependence, environmental variability, and demographic stochasticity from population counts: An example using Wytham Wood great tits. In Population Biology of Passerine Birds, pages 89–102. Springer, Berlin, Heidelberg.

Lebreton, J. D. (2005). Age, stages, and the role of generation time in matrix models. Ecological Modelling, 188(1):22–29.

Lebreton, J.-D. and Gonzáles-Dávila, G. (1993). An introduction to models of subdivided populations. Journal of Biological Systems, 01(04):389–423.

Lotka, A. J. (1939). Théorie analytique des associations biologiques. Hermann et Cie, Paris.

Luo, J. and Fox, B. J. (1990). Life-table comparisons between two ground squirrels. Journal of Mammalogy, 71(3):364–370.

Meyer, C. D. (2000). Matrix analysis and applied linear algebra, volume 71. SIAM.

Nerman, O. and Jagers, P. (1984). The stable doubly infinite pedigree process of supercritical branching populations. Zeitschrift fuer Wahrscheinlichkeitstheorie und Verwandte Gebiete, 65(3):445–460.

Ozgul, A., Oli, M. K., Armitage, K. B., Blumstein, D. T., and Van Vuren, D. H. (2009). Influence of local demography on asymptotic and transient dynamics of a yellow-bellied marmot metapopulation. American Naturalist, 173(4):517–530.

Pavard, S. and Coste, C. F. D. (2020). Goodman, Keyfitz & Pullum (1974) and the population frequencies of kinship relationships. Theoretical Population Biology, 133:15–16.

Pullum, T. W. (1982). The eventual frequencies of kin in a stable population. Demography, 19(4):549–565.

Rees, M., Childs, D. Z., and Ellner, S. P. (2014). Building integral projection models: A user’s guide. Journal of Animal Ecology, 83(3):528–545.

Roth, G. and Caswell, H. (2016). Hyperstate matrix models: extending demographic state spaces to higher dimensions. Methods in Ecology and Evolution, 7(12):1438–1450.

Tuljapurkar, S. (1993). Entropy and convergence in dynamics and demography. Journal of Mathematical Biology, 31(3):253–271.

Tuljapurkar, S., Zuo, W., Coulson, T., Horvitz, C., and Gaillard, J. M. (2020). Skewed distributions of lifetime reproductive success: beyond mean and variance. Ecology Letters, 23(4):ele.13467.

Tuljapurkar, S., Zuo, W., Coulson, T., Horvitz, C., and Gaillard, J. M. (2021). Distributions of LRS in varying environments. Ecology Letters, pages 1328–1340.

Tuljapurkar, S. D. (1982). Population Dynamics in Variable Environments. IV. Weak ergodicity in the Lotka Equation. Journal of Mathematical Biology, 14:221–230.

Varga, R. S. (2000). Matrix Iterative Analysis, volume 27 of Springer Series in Computational Mathematics. Springer, Berlin, Heidelberg.

Waugh, W. A. O. (1981). Application of the Galton-Watson process to the kin number problem. Advances in Applied Probability, 13(4):631–649.

Young, K. E. and Keith, E. O. (2011). A comparative analysis of cetacean vital rates using matrix population modeling analysis of cetacean vital rates. International Journal of Applied Science and Technology, 1(6):261–277.

